# Single-cell membrane potential fluctuations evince network scale-freeness and quasicriticality

**DOI:** 10.1101/498477

**Authors:** James K Johnson, Nathaniel C. Wright, Ji Xia, Ralf Wessel

## Abstract

What information single neurons receive about general neural circuit activity is a fundamental question for neuroscience. Somatic membrane potential fluctuations are driven by the convergence of synaptic inputs from a diverse cross section of upstream neurons. Furthermore, neural activity is often scale-free implying that some measurements should be the same, whether taken at large or small scales. Together, convergence and scale-freeness support the hypothesis that single membrane potential recordings carry useful information about high-dimensional cortical activity. Conveniently, the theory of “critical branching networks” (a purported explanation for scale-freeness) provides testable predictions about scale-free measurements which are readily applied to membrane potential fluctuations. To investigate, we obtained whole-cell current clamp recordings of pyramidal neurons in visual cortex of turtles with unknown genders. We isolated fluctuations in membrane potential below the firing threshold and analyzed them by adapting the definition of “neuronal avalanches” (spurts of population spiking). The membrane potential fluctuations we analyzed were scale-free and consistent with critical branching. These findings recapitulated results from large-scale cortical population data obtained separately in complementary experiments using microelectrode arrays (previously published (Shew et al., 2015)). Simultaneously recorded single-unit local field potential did not provide a good match; demonstrating the specific utility of membrane potential. Modeling shows that estimation of dynamical network properties from neuronal inputs is most accurate when networks are structured as critical branching networks. In conclusion, these findings extend evidence for critical branching while also establishing subthreshold pyramidal neuron membrane potential fluctuations as an informative gauge of high-dimensional cortical population activity.

**Significance Statement:** The relationship between membrane potential dynamics of single neurons and population dynamics is indispensable to understanding cortical circuits. Just as important to the biophysics of computation are emergent properties such as scale-freeness, where critical branching networks offer insight. This report makes progress on both fronts by comparing statistics from single-neuron whole-cell recordings to population statistics obtained with microelectrode arrays. Not only are fluctuations of somatic membrane potential scale-free, they match fluctuations of population activity. Thus, our results demonstrate appropriation of the brain’s own subsampling method (convergence of synaptic inputs), while extending the range of fundamental evidence for critical branching in neural systems from the previously observed mesoscale (fMRI, LFP, population spiking) to the microscale, namely, membrane potential fluctuations.

## Introduction

How do cortical population dynamics impact single neurons? What can we learn about cortical population dynamics from single neurons? These questions are central to neuroscience. Uncovering the functional significance of multiscale organization within cerebral cortex requires knowing the relationship between the dynamics of networks and individual neurons within them (Nunez et al., 2013).

For pyramidal neurons in the visual cortex, somatic spike generation is ambiguously related to presynaptic firing (Tsodyks and Markram, 1997; Brunel et al., 2014; Gatys et al., 2015; Stuart and Spruston, 2015; Moore et al., 2017). Such neurons pass spiking information to many postsynaptic neurons (Lee et al., 2016). However, a presynaptic pool with multifarious neighboring and distant neurons (Hellwig, 2000; Wertz et al., 2015) provides excitatory and inhibitory synaptic inputs throughout the soma and complex dendritic architecture (Magee, 2000; Larkum et al., 2008; Moore et al., 2017). Input propagation to the axon hillock has both active and passive features (London and Hausser, 2005), and the membrane potential (V_m_) response is increasingly non-linear near the action potential threshold. Thus, such details of network propagation give membrane potential more utility than focusing solely on spiking.

Most computational neuroscientists focus on spiking data because spikes are the “currency of the brain” (Wolfe et al., 2010), and recording extracellular activity is more straightforward than obtaining whole-cell recordings. Yet, the paucity of single-neuron spiking (Shoham et al., 2006), and lack of foreknowledge about connections (Helmstaedter, 2013) makes extracellular single-unit observation an impoverished source of information about neuronal circuits. In contrast, subthreshold V_m_ fluctuations contain rich information about the circuits containing each neuron (Sachidhanandam et al., 2013; Petersen, 2017). Integral to gaining a neuron’s view of the brain is uncovering relationships between the statistics of V_m_ fluctuations and fluctuations of local spiking, and then contrasting against other plausible one-dimensional signals.

We look for such relationships in the strict predictions and rigorous measurements of scale-freeness used to identify a fragile network connectivity pattern known as “critical branching”. This pattern confers emergent properties valuable for information processing, such as higher susceptibility and dynamic range (Haldeman and Beggs, 2005; Beggs, 2007; Shew and Plenz, 2013; Shriki and Yellin, 2016; Timme et al., 2016). The pattern is as follows: on average over all neuronal avalanches (fluctuations of spiking above baseline (Friedman et al., 2012)), one spike leads to *exactly* one other spike. In most arbitrary networks there is less or more than one; these are “subcritical” and “supercritical” respectively. Among the dazzling emergent properties of “criticality” are universality, self-similarity, and scale-free correlations (Stanley, 1999).

These are as follows: A “universality class” refers to a set of incongruous systems exhibiting identical statistics only at their “critical points”. Self-similarity includes power-laws in geometrical analysis of avalanches (power-laws are “scale-invariant”, popularly called “scale-free”). Avalanches of any duration have the same average shape if normalized (Shaukat and Thivierge, 2016). Their area grows with duration as a power-law (Sethna et al., 2001). However, the observation tool must be consistent with event propagation (Priesemann et al., 2009; Yu et al., 2014; Levina and Priesemann, 2017). Additionally, correlations are scale-free, meaning correlation length and time are infinite (Chialvo, 2010). Any input has a nonzero chance of propagating forever or to every point.

In summary, the theory of critical branching networks offers superb standards of comparison for three reasons: neuronal avalanche analysis applies to membrane potentials, it offers promising insights, and makes precise predictions about the statistics of fluctuation geometry. We can study both V_m_ fluctuations and criticality with one simple question: Do V_m_ fluctuations match the scale-free statistics of cortical populations (**Figure 1**)?

**Figure 1:**
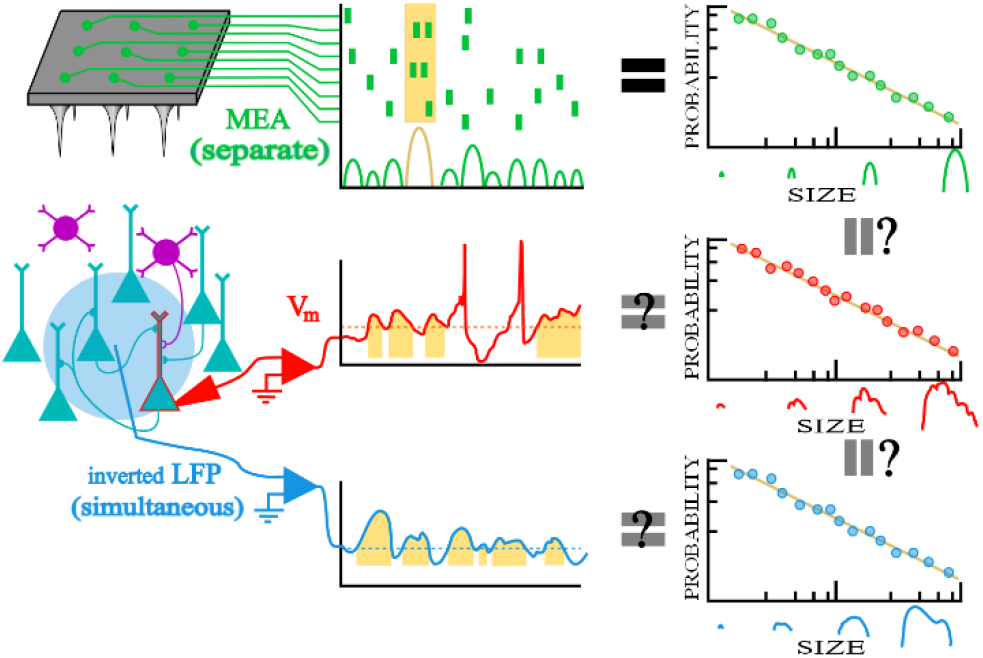
Will fluctuations in somatic membrane potential and comparable signals reflect the scale-free nature of neuronal avalanches from microelectrode array data? A recurrent network with excitatory (teal) and inhibitory (purple) neurons is measured in three ways: microelectrode array (MEA)(green/upper), whole-cell recording (red/middle), LFP (blue/bottom). Neuronal avalanches (highlighted in gold) are inferred from the population raster and fluctuations are analyzed like avalanches for the V_m_ and inverted LFP signals. Neuronal avalanches are defined as spurts of activity with quiet periods between them for MEA or excursions above the 25^th^ percentile for continuous non-zero data. The ultimate question is whether membrane potential fluctuations will recapitulate the entire neuronal avalanche analysis previously conducted on MEA data, including power-laws in size and duration as well as a universal avalanche shape. This is abridged in the right most column which illustrates power-law distributions.

To address this question, we simultaneously recorded somatic V_m_ from pyramidal neurons and local field potential (LFP) in visual cortex and performed avalanche analysis on fluctuations. We found that subthreshold V_m_ fluctuation statistics match published microelectrode array (MEA) data. We used surrogate testing to show why negative LFP fluctuations don’t match and modeling to demonstrate dependence on critical branching.

## Methods

### Surgery and Visual Cortex

All procedures were approved by Washington University’s Institutional Animal Care and Use Committees and conform to the guidelines of the National Institutes of Health on the Care and Use of Laboratory Animals. Fourteen adult red-eared sliders (*Trachemys scripta elegans*, 150-1000 g) were used for this study, their genders were not recorded. Turtles were anesthetized with Propofol (2 mg Propofol/kg), then decapitated. Dissection proceeded as described previously (Saha et al., 2011; Crockett et al., 2015; Wright et al., 2017a).

To summarize, immediately after decapitation, the brain was excised from the skull, with right eye intact, and bathed in cold extracellular saline (in mM, 85 NaCl, 2 KCl, 2 MgCl_2_*6H_2_O, 20 Dextrose, 3 CaCl_2_-2H_2_O, 45 NaHCO_3_). The dura was removed from the left cortex and right optic nerve, and the right eye hemisected to expose the retina. The rostral tip of the olfactory bulb was removed, exposing the ventricle that spans the olfactory bulb and cortex. A cut was made along the midline from the rostral end of the remaining olfactory bulb to the caudal end of the cortex. The preparation was then transferred to a perfusion chamber (Warner RC-27LD recording chamber mounted to PM-7D platform) and placed directly on a glass coverslip surrounded by Sylgard. A final cut was made to the cortex (orthogonal to the previous and stopping short of the border between medial and lateral cortex) allowing the cortex to be pinned flat, with ventricular surface exposed. Multiple perfusion lines delivered extracellular saline to the brain and retina in the recording chamber (adjusted to pH 7.4 at room temperature).

We used a phenomenological approach to identify the visual cortex, described previously (Shew et al., 2015). In general, this region was centered on the anterior lateral cortex, in agreement with voltage-sensitive dye studies (Senseman and Robbins, 1999; Senseman and Robbins, 2002). Anatomical studies identify this as a region of cortex receiving projections from lateral geniculate nucleus (Mulligan and Ulinski, 1990).

### Intracellular Recordings

For whole-cell current clamp recordings, patch pipettes (4-8 MΩ) were pulled from borosilicate glass and filled with a standard electrode solution (in mM; 124 KMeSO_4_, 2.3 CaCl_2_-2H_2_O, 1.2 MgCl_2_, 10 HEPES, 5 EGTA) adjusted to pH 7.4 at room temperature. Cells were targeted for patching using a differential interference contrast microscope (Olympus). Membrane potential recordings were collected using an Axoclamp 900A amplifier, digitized by a data acquisition panel (National Instruments PCIe-6321), and recorded using a custom LabVIEW program (National Instruments), sampling at 10 kHz. We excluded cells that did not display stable resting membrane potentials for long enough to gather enough avalanches. Up to 3 whole-cell recordings were made simultaneously. In total, we obtained recordings from 51 neurons from 14 turtles.

Recorded V_m_ fluctuations taken in the dark (no visual stimulation) were interpreted as ongoing activity. Such ongoing cortical activity was interrupted by visual stimulation of the retina with whole-field flashes and naturalistic movies as described previously (Wright et al., 2017a; Wright et al., 2017b; Wright and Wessel, 2017). An uninterrupted recording of ongoing activity lasted for 2 to 5 minutes.

A sine-wave removal algorithm was used to remove 60 Hz line noise. Action potentials in turtle cortical pyramidal neurons are relatively rare. An algorithm was used to detect spikes, the V_m_ recordings between spikes were extracted and filtered from 0 to 100 Hz. Membrane potential recordings were de-trended by subtracting the 5^th^ percentile in a sliding 2 s window. The resulting signal was then shifted to have the same mean value as before subtraction. De-trending did not affect the size of membrane potential fluctuations (data not shown).

### Extracellular Recordings

Extracellular recordings were achieved with tungsten microelectrodes (microprobes heat-treated tapered tip), with approximately 0.5 MΩ impedance. Electrodes were slowly advanced through tissue under visual guidance using a manipulator (Narishige), while monitoring for activity using custom acquisition software (National Instruments). The extracellular recording electrode was located within approximately 300 μm of patched neurons. Extracellular activity was collected using an A-M Systems Model 1800 amplifier, band-pass filtered between 1 Hz and 20,000 Hz, digitized (NI PCIe-6231), and processed using custom software (National Instruments). Extracellular recordings were down-sampled to 10,000 Hz and then filtered (100 Hz low-pass), yielding the local field potential (LFP). The LFP was filtered and detrended as described above (see Intracellular Recordings), except that the mean of the entire signal was subtracted, and the signal was multiplied by −1 before it was detrended. This final inverted signal is commonly featured in literature as negative LFP or nLFP (Kelly et al., 2010; Kajikawa and Schroeder, 2011; Okun et al., 2015; Ness et al., 2016).

### Experimental Design and Statistical Analysis

#### Set-wise comparisons

In order to measure differences between sets of statistics we rely on three non-parametric measures. We use the MATLAB Statistics and Machine Learning Toolbox implementation of Fisher’s exact test (Hammond et al., 2015). This lets us measure the effect size (Odds Ratio *r_OR_*) and statistical significance (p value) of finding that consistency with criticality is more frequent or less frequent in an experimental group than a control group.

To quantify the similarity between the exponents measured in different sets of data we use the MATLAB Statistics and Machine Learning Toolbox implementations of the *exact* Wilcoxon rank sum test (Hammond et al., 2015) and the *exact* Wilcoxon signed rank test. In both cases effect size, *r_SDF_* is measured by the simple difference formula (Kerby, 2014). The rank sum test is used when comparing non-simultaneous recordings, such as comparing MEA data with V_m_ data. The signed rank test is used when comparing data that can be paired, such as V_m_ data to concurrent LFP. When comparing whether a dataset differs from a specific value, we can use the sign test.

The significance level is set at p=0.05 for all tests; we are not making multiple comparisons (Bender and Lange, 2001).

#### Random surrogate testing

It is possible that scale-free observations have an origin in independent random processes of a kind previously demonstrated (Touboul and Destexhe, 2017). To control for this, we phase-shuffled the V_m_ fluctuations using the amplitude adjusted Fourier transform (AAFT) algorithm (Theiler, 1992). This tests against the null hypothesis that a measure on a time series can be reproduced by performing a non-linear rescaling of a linear Gaussian process with the same autocorrelation (same Fourier amplitudes) as the original process. Phase information is randomized, which removes higher-order correlations but preserves the scale-free power-spectrum.

The AAFT tests only higher-order correlations, but a simpler algorithm tests against the null hypothesis that an *un-rescaled* linear Gaussian process with the same autocorrelation as the original process can produce the same results (Theiler, 1992). This is known as the Unwindowed Fourier Transform (UFT). Once we see what measures depend on the higher-order correlations with the AAFT we can use the UFT to see how measures depend on the non-Gaussianity (non-linear rescaling) which is inherent to excitable membranes. Using the UFT alone would make it difficult to attribute whether statistically significant differences are due to the rescaling or to the higher-order correlations (Rapp et al., 1994).

We performed AAFT and UFT on each V_m_ time series once, and then compared how the two datasets performed on every metric used in this study. The datasets were compared with a matched Wilcoxon sign rank test implemented via MATLAB’s statistics tool box. Doing the comparison at a dataset level allowed us to obtain a discrimination statistic for every metric we used without repeating the computationally expensive analysis procedure hundreds or thousands of times on every V_m_ trace. With enough individual recordings in each dataset the matched Wilcoxon sign rank test is a reliable measure, which empowered us to efficiently compare all important metrics.

#### Neuronal avalanche analysis

Neuronal avalanches were defined by methods analogous to (Poil et al., 2012), which are used for uninterrupted ongoing signals whereas methods based on event detection (Beggs and Plenz, 2003) require periods of non-activity. A threshold is defined, and an avalanche starts when the signal crosses the threshold from below and ends when the signal crosses the threshold from above. The choice of threshold is a free parameter and we set it to the 25^*th*^ percentile. Several nearby percentiles were tested and gave similar power-law exponents between the 15^th^ to 50^th^ percentile. However, the number of avalanches is much less with either extreme. The 25^th^ percentile was chosen because it gave many avalanches and changing the threshold to maximize the number of avalanches for individual trials adds complexity and opportunity for artifacts.

We quantified each neuronal avalanche by its size *A* and its duration *D*. The avalanche size is the area between the processed V_m_ recording and the baseline. The baseline is another free parameter that was set at the second percentile of the processed V_m_ recording. The second percentile was chosen because its value is more stable than the absolute minimum. The avalanche duration *D* is the time between threshold crossings.

The lower limit of avalanche duration is defined by the membrane time constant which has been reported to be between 50 and 140 ms for the turtle brain at room temperature (Ulinski, 1990; Larkum et al., 2008). We took a conservative approach by setting the limit at less than half the lower bound on membrane time constant which was significantly less than the lower cut-off from power-law fits. Only avalanches of duration larger than 20 ms were included in the analysis. Thus, we avoided artificially retaining only the events most likely to be power-law distributed.

Following the procedure described above, each processed V_m_ recording of uninterrupted ongoing activity (i.e., a recording of 2 to 5 minutes duration) yielded 327 ± 148 (mean ± standard deviation) avalanches. This is insufficient for rigorous statistical fitting on recordings individually (Clauset et al., 2009). Therefore, we grouped avalanches from multiple recordings of ongoing activity of the same cells. Each cell produced between 3 and 19 recordings of ongoing activity (2 to 5 minutes duration each recording), with trials recorded intermittently over a period of 10 to 60 minutes. We grouped recordings based on whether they occurred in the first or second 20-minute period since the beginning of recording from that neuron. Then all the avalanches from the first or second 20-minute period were grouped together with one data object (the group) storing the size, and duration of each avalanche. It is rare for neurons to have recordings in the third 20-minute periods, so this data was not included. Since there was a slow drift in the mean membrane potential over a period of several minutes, we scaled the avalanche sizes from each recording to have the same median as other recordings from the same group. On average 4 recordings were possible in each 20-minute period. There were 51 neurons with multiple recordings of ongoing activity in the first 20-minutes of experimentation (thus 51 recording groups). Of these, 18 neurons had an additional 20-minute period with more than one recording. This produced a total of 69 groups with 1346 ± 1018 (mean ± standard deviation) avalanches for each group. Of these 69 groups, 57% had more than 1000 avalanches. The largest number of avalanches was 7495 and the smallest was 313. Only 5 groups had less than 500 avalanches. We report on the 51 groups from the first 20-minute period separately from the 18 groups with recordings from the second 20-minute period of experimentation.

For each group, we evaluated the avalanche size and duration distributions with respect to power laws. To test whether a distribution followed a power law, we applied the rigorous statistical fitting routine described previously (Clauset et al., 2009). We tested three power-law forms: *P*(*x*) ∝ *x^−α^* (with and without truncation) (Deluca and Corral, 2013), as well as a power-law with exponential cut-off *P*(*x*) ∝ *x^−α^e*^−*x*/*r*^. We compared these against lognormal and exponential alternative (non-power-law) hypotheses. Distribution parameters were estimated using Maximum Likelihood Estimation (MLE) and the best model out of those fitted to the data was chosen using the Akaike Information Criterion (Bozdogan, 1987). It should be acknowledged that a small power-law region in the truncated form would be suspect for false positives, likewise for a strong exponential cut-off (Deluca and Corral, 2013). Finally, to decide whether a fitted model was plausible, pseudo-random datasets were drawn from a distribution with the estimated parameters and then the fraction which had a lower fit quality (Kolmogorov-Smirnov distance) than the experimental data was calculated. If this fraction, called the comparison quotient *q*, was greater than 0.10, the best fit model (according to the Akaike Information Criterion) was accepted as the best candidate. Otherwise, the next best model was considered.

We applied several additional steps and strict criteria to control for false positives. One such step was assessing whether the scaling relation was obeyed over the whole avalanche distribution for each group (not just the portion above the apparent onset of power-law behavior). The scaling relation is another power-law 〈*A*〉(*D*) ∝ *D^γ^* predicting how the measured size of avalanches increase geometrically with increasing duration (on average). For any data set which has three power-laws, 〈*A*〉(*D*) ∝ *D^γ^* (scaling relation), *P*(*A*) ∝ *A*^−*τ*^ (size distribution), and *P*(*D*) ∝ *D^−β^* (duration distribution), the scaling relation exponent is predicted by the other two exponents by 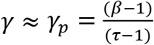 (Scarpetta et al., 2018). Note that *γ_p_* = 1 is a trivial value because it implies 〈*A*〉(*D*) ∝ *D* and that would suggest individual avalanches were just noise symmetric about a constant value. This would mean that the average avalanche shape is just a flat line at some constant of proportionality, 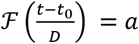, where 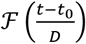 is a function describing the shape of an avalanche of duration *D* and *t*_0_ is the beginning of the avalanche and *a* is a constant.

##### Standards for consistency with critical point behavior

We applied four standardized criteria to provide a transparent and systematic way to produce a binary classification, either “no inconsistencies with activity near a critical point were detected” or “some inconsistencies with activity near a critical point were detected”.

First, a collection of avalanches must be power-law distributed in both its size and duration distributions.

Second, the collection of avalanches must have a power-law scaling relation as determined by *R*^2^ > 0.95 (coefficient of determination) for linear least squares regression to a log-log plot of average size vs durations: log(〈*A*〉(*D*)) *~γ* log(*D*) + *b*. This *R*^2^ represents the best that any linear fit can achieve and must include all the avalanches, not a subset. We denote the scaling exponent (slope from linear regression) from this fit as *γ_f_*.

Third, the scaling relation exponent predicted by theory (denoted as *γ_p_*) must correspond to a trendline on a log-log scatter plot of 〈*A*〉(*D*) whose *R*^2^ is within 90% of the best-case fitted trendline from the second criterion. Again, the *R*^2^ for the predicted scaling relation is calculated across all avalanches, and not just the subset above the inferred lower cut-off of power-law behavior (which was found for the first criterion). This cross-validates agreement with theory.

Fourth, the fitted scaling relation exponent must be significantly greater than 1: (*γ_f_* − 1) > *σ_γ_f__* where *σ_yf_* is the standard error. This last requirement eliminates scaling that might be trivial in origin. It is measured after getting the fitted scaling relation exponent for all the data so that a dataset standard deviation can be determined. It is necessary to also check that the set of scaling relation exponents from the power-law fits to all avalanche sets is significantly different from 1 at a dataset level. A scaling relation exponent equal to one suggests a linear relationship between mean-size and duration which is not consistent with criticality in neural systems (Haldeman and Beggs, 2005).

Our four-criterion test cannot measure distance from a critical point nor eliminate all risk of false positives. To complete our analysis, we also look at three additional factors, whether exponent values match exponent values from *other* experiments as expected from the universality prediction of theory, whether all the exponents within our data set have similar scaling relation predictions, and lastly whether the avalanches within our data set exhibit shape collapse across all the recordings.

##### Applying shape collapse, quantitative and qualitative analysis

Shape collapse is a very literal manifestation of scale-invariance (also called “self-similarity”)(Sethna et al., 2001; Beggs and Plenz, 2003; Friedman et al., 2012; Pruessner, 2012; Timme et al., 2016). Avalanches of different durations should rise and fall in the same way on average. This average avalanche profile is called a scaling function. The average avalanche profile for avalanches of duration *D* is predicted to be 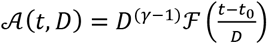 where *D^(γ−1)^* is the power-law scaling coefficient which modulates the height of the profile and 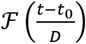 is the universal scaling function itself (normalized in time). Shape collapse analysis provides an independent estimate of the scaling relation exponent *γ_SC_*, which is only expected to be accurate at criticality (Sethna et al., 2001; Scarpetta and de Candia, 2013; Shaukat and Thivierge, 2016), and a visual test of conformation to an empirical scaling function.

Exponent estimation is very sensitive to the unrelated, intermediate rescaling steps involved in combining the avalanches from multiple recordings into one group. To get an estimate of the scaling relation exponent for each group, *γ_SC_*, we average the scaling exponents *γ_i_* found individually for each recording in that group (*i* denotes the *i*^th^ recording, SC for “shape collapse”).

Naturally, individual avalanche profiles are vectors of variable length *D*. We must first “rescale in time” to make them vectors of equal length without losing track of what each vector’s original duration was. We do that by linearly interpolation with 20 evenly spaced points. So, the *j*^th^ avalanche profile of the *i*^th^ recording is denoted as a 20-element vector 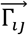, (where the top arrow denotes a vector).

Next, the set of all profiles from recording *i* with the *exact same duration D*, denoted as **Γ**_**D**_*i*__ where bold indicates a set, were averaged and divided by a test scaling factor 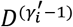. We define this as 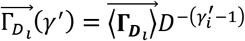. The prime indicates a test rescaling. The average is over all vectors in the set **Γ**_**D**_*i*__. The choice of *γ_i_* was optimized using MATLAB’s fminsearch function to minimize the mean relative error between the average over all durations 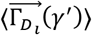 and the set members 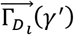 so that for recording *i*:

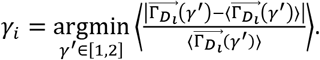

This error minimization and applying the rescaling is the “collapse” in “shape-collapse”.

Once we have the *γ_i_* for the avalanches in each individual recording of ongoing activity we compare the average, *γ_SC_* = 〈γ_i_〉, to the predicted and fitted scaling relation exponents for the group of recordings, *γ_p_* and *γ_f_* (statistical comparison tests are described in a previous section). Thus, quantitative analysis of shape collapse was done by comparing *γ_SC_*, *γ_p_*, and *γ_f_* for each of the 69 groups individually.

Visual assessment of how well avalanche profiles can be described by one universal scaling function, 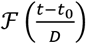 supports the quantitative exponent estimation. This was carried out by averaging all the profiles within specific *duration bins* (regardless of trial or group) and plotting them on top of one another. A very large number of avalanches are needed so we combine avalanches from all 69 groups. However, the resting membrane potential differs from recording to recording and cell to cell. Therefore, avalanche profiles from different recordings are vertically misaligned. To combine avalanches profiles from different recordings we divided all the profiles by a scalar value unique to each recording: the time average over all the collapsed profiles. This produce rescaled and mean-shifted profiles (double prime) 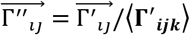 (where *k* ∈ [1,20] denotes the interpolated time point). The set of avalanches from each recording were thus aligned, but individual variability was preserved and thus profiles from different recordings could be averaged without introducing artifacts. This set, **Γ″_ij_** contained a total of 106,220 shifted and rescaled profiles for the V_m_ data.

The set of shifted and rescaled profiles falling into a duration bin is denoted 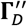. Each duration bin then provides its own estimate of the scaling function 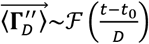. For each bin, *D* was defined as the average duration of all constituent profiles. If less than 700 avalanches had a particular duration, we included the next longest duration iteratively until we met or exceeded 700 avalanches. This only applied to long durations. The choice of 700 was made because it allowed us smooth averaging and without excessively wide duration bin widths.

We also assessed the mean curvature of avalanche profiles from the rescaled profile for a particular duration 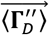. This allows us to plot how curvature depends on duration. Mean curvature 〈κ〉 is defined like so (*k* still denotes time points):

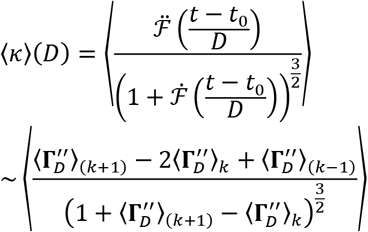

### Model Simulations

We simulated a model network consisting of *N* = 10^4^ binary probabilistic model neurons. The model neurons form a directed random network (Erdős–Rényi random graph), where the probability that neuron *j* connects to neuron *i* is *c*. In a network of *N* neurons, this results in a mean in-degree and out-degree of *cN*. We tested nine not quite evenly distributed values of connection probabilities *c* ∈ [0.5,1,3,5,7.5,10,15,20,25] × 10^−2^.

The strength of the connection from neuron *j* to neuron *i* is quantified in terms of the network adjacency or weight matrix *W* with the fortune of having a simple and intuitive meaning. For each existing connection from neuron *j* to neuron *i, W_ij_* is the direct change in the probability that neuron *ı* will fire at the next timestep if neuron *j* spikes in the current time step.

The dynamics of this network is well-characterized by the largest eigenvalue *λ* of the network weight matrix *W*, with criticality occurring at *λ* = 1 (Kinouchi and Copelli, 2006; Larremore et al., 2011b; Larremore et al., 2011a; Larremore et al., 2014). The physical interpretation of *λ* is a “branching parameter”(Haldeman and Beggs, 2005) that governs expected number of spikes immediately caused by the firing of one neuron. If *λ* = 1 then one spike causes one other spike on average, while if *λ* > 1 one spike causes more than one on average and vice versa.

We tested five different values of largest eigenvalue at, near and far from criticality *λ* ∈ [0.9,0.95,1,1.015,1.03]. A fraction *χ* of the neurons are designated as inhibitory. This is done by multiplying all outgoing connections of an inhibitory neuron by −1. We tested nine different values of the fraction of inhibitory neurons in the range from 0 to 0.25, thus including the value 0.2, corresponding to the fraction of inhibitory neurons in the mammalian cortex (Meinecke and Peters, 1987). The magnitudes of non-zero weights are independently drawn from a distribution of positive numbers with mean *η*, where the distribution is uniform on [0,2*η*], and *η* is given by *η* = *λ*/(*cN*(1 − 2χ)). The maximum eigenvalue is then fine-tuned by dividing *W* by the current maximum eigenvalue and set to the exactly desired value *W* = *λW′*/*λ* where *W′* and *λ*^1^ are the matrices and eigenvalues before correction.

The binary state *S_i_*(*t*) of neuron *i* at time *t* denotes whether the model neuron spikes (*S_i_*(*t*) = 1) or does not spike (*S_i_*(*t*) = 0) at time *t*. At each time step, the states of all neurons are updated synchronously according to the following update rule:

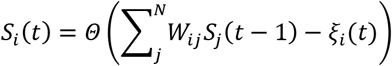

where *ξ_i_*(*t*) is a random number on [0 1] drawn from a uniform distribution, and *Θ* is the Heaviside step function. In addition to this update rule, a refractory period of 2 time-steps (translated to approximately 4 ms) was imposed for certain parameter conditions. A simulation begins with initiating the activity of one randomly-chosen excitatory neuron and continuing the simulation until overall network activity had ceased. The process was then repeated.

From the simulated binary states of 10^4^ model neurons, we extracted three measures of simulated activity. First, the network activity 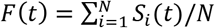 is the fraction of neurons spiking at time *t*. Second, the input to model neuron *i* at time *t* is 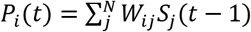, which is almost always positive for our parameters. Note that 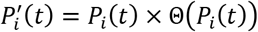 directly represents the probability for the neuron to spike at time *t*. Third, we constructed a proxy for the V_m_ signal, Φ_*i*_(*t*) = (*α_h_* * *P_i_*)(*t*), by convolving the input *P_i_*(*t*) with an alpha function: 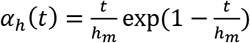 with *h_m_* = 2 time steps (assumed to be about 4 *ms*).

A total of 405 different parameter combinations (connection density, inhibition, maximum eigenvalue) were simulated. Each combination was simulated 10 times. Based on the connection probability *c* and the fraction of inhibition *χ*, we distinguish four regions in parameter space classified according to the behavior of the critical model, i.e., *λ* = 1.

The first region is the “positive weights” region. Without inhibition activity increases or dies out in accordance with the branching parameter. This region is defined by *χ* = 0. With moderate inhibition and dense connectivity there is a region of parameter space we call “quiet”; activity lasts only slightly longer than in a system with no inhibition. This region is defined by the ex-post-facto boundaries *c* ≥ *e*^11*χ*^/25 and *χ* > 0. Further increasing inhibition relative to connection density produces a behavior like “up and down” states (or “telegraph noise”) (Sachdev et al., 2004; Millman et al., 2010). We call this the “switching” regime because network activity switches between a low mean and a high mean. This region is defined by < *e*^11*χ*^/25, and *c* ≥ (10*e*^12*χ*^ − 13)/100 and *χ* > 0. When inhibition is high relative to connection density the system enters the “ceaseless” region where stimulating one neuron causes activity that effectively never dies out.

We set the refractory period to two time-steps if the network is in the ceaseless regime or if it is in the switching regime and spends greater than 50% of its time about a high mean (an “up state”). This is determined by an initial testing cycle before simulation begins.

We performed avalanche analysis on each of the simulated signals using the methods described above for membrane potential recordings. If the network is in the switching regime, we only perform analysis on the periods when the network is in the mode (high or low mean) in which it spends the majority of its time. As before, the 25th percentile defined the avalanche threshold. If the signal had negative values, as in the case of single neuron V_m_ proxies in networks with inhibition, the signal was shifted by subtracting the 2nd percentile. To obtain good statistics, we continued stimulating and extracting avalanches until a simulation either reached 10^4^ avalanches, or 5 × 10^3^ avalanches and a very large file size or a very long computational time. This ensured there were between two and ten thousand avalanches per trial.

## Results

Single-neuron membrane potential (V_m_) fluctuations are thought to be dominated by synaptic inputs from multitudes of presynaptic neurons (Stepanyants et al., 2002; Brunel et al., 2014; Petersen, 2017). It is crucial that neuroscience gain a thorough understanding of the relationship between subthreshold V_m_ fluctuations and population activity. A basic step is to compare statistical analyses, especially analyses where a meaningful relationship is expected. We asked whether an avalanche analysis on V_m_ fluctuations would reveal the same signatures of scale-freeness and critical network dynamics found in measures of population activity (**Figure 1**) (Friedman et al., 2012; Shew et al., 2015; Marshall et al., 2016). To address this comparison across organizational levels, we recorded V_m_ fluctuations from 51 pyramidal neurons in visual cortex of 14 turtles and assessed evidence for critical network dynamics from these recordings.

In a model investigation we corroborated results evaluated the conditions needed to enable inferring dynamical network properties from the inputs to single neurons. Finally, we extended the analysis to other commonly recorded time series of neural activity for comparison with the information content of V_m_ fluctuations about the dynamical network properties.

### Membrane Potential Fluctuations Reveal Signatures of Critical Point Dynamics

We obtained whole-cell recordings from pyramidal neurons in the visual cortex of the turtle ex-vivo eye-attached whole-brain preparation (**Figure 2A**). Recorded V_m_ fluctuations taken in the dark (no visual stimulation) were interpreted as ongoing activity. We analyzed the recorded ongoing V_m_ fluctuations employing the concept of “neuronal avalanches” (Beggs and Plenz, 2003; Poil et al., 2012; Shew et al., 2015), which are positive fluctuations of network activity. For continuous time-series such as the V_m_ recording, one selects a threshold and a baseline. We defined a neuronal avalanche based on the positive threshold crossing followed by a negative threshold crossing of the V_m_ time series (Poil et al., 2012; Hartley et al., 2014; Larremore et al., 2014; Karimipanah et al., 2017b). We quantified each neuronal avalanche by (i) its size A, i.e., the area between the curve and the baseline, and (ii) its duration *D*, i.e., the time between threshold crossings (**Figure 2B**).

**Figure 2:**
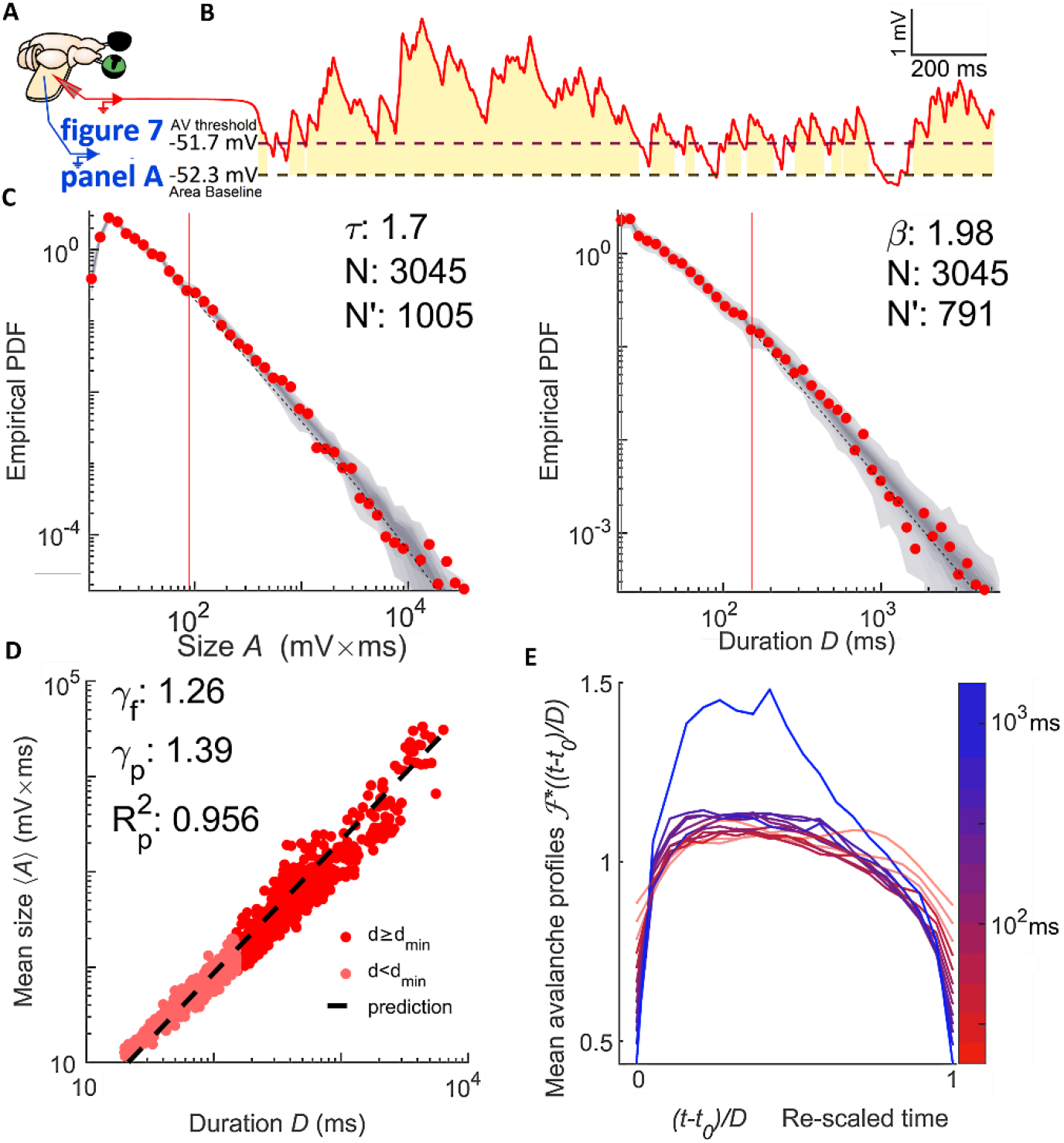
Membrane potential fluctuations reveal signatures of critical point dynamics. **Panel A** shows the whole-brain eye attached joint V_m_ and LFP recording preparation. **Panel B** shows that the membrane potential (red) is thresholded at the 25^th^ percentile (a dashed line). Avalanches are defined by excursions above this threshold. The gold region represents the size of the avalanche, which is the area between the signal and its 2^nd^ percentile (a dashed line). The duration of the avalanche is the duration of the excursion. **Panel C** shows the size (left) and duration (right) distributions of V_m_ inferred avalanches when data is combined from seven recordings from the same neuron falling in the same 20-minute period. The comparison quotients (q) are both above 0.10 (0.878 and 0.874 respectively), indicating that the size and duration distributions were better fits to power-laws at the given cut-off than 87% of power-laws produced by a random number generator with the same parameters (shown as a grey density cloud). N’ indicates the number of avalanches above the lower cut-off of the fit (red vertical line) and N indicates the total number of avalanches. Size duration exponent denoted with *τ* while *β* is used for duration. **Panel D** shows the scaling relation which is a function relating average avalanche size to each given duration. The predicted exponent (*γ_p_*) successfully explains 95.6% of the variance of a log-log representation of the data. A linear least squares regression could explain 96.7% and gives the fitted exponent (*γ_f_*). Therefore, *γ_p_* comes within 1.2% of the best linear explanation despite a 10% difference in exponent values. **Panel E** shows shape collapse. Each line represents the average time-course of an avalanches of a given duration. The color indicates the duration according to the scale bar. Durations below 50 ms (the lower bound on turtle pyramidal time-constants) are made translucent and slightly thickened. This shape collapse represents the global collapse across all recordings in all cells. This confirms that a universal scaling function, 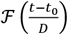, is present. For the seven recordings in the group represented in panels C & D, the mean scaling relation exponent derived from shape collapse was *γ_SC_* = 1.23 a disagreement of 2.2% relative to *γ_f_*.

To quantify the statistics of avalanche properties, we applied concepts and notations from the field of “critical phenomena” in statistical physics (Nishimori and Ortiz, 2011; Pruessner, 2012). Because the critical point of a critical branching network is such a small target for any naturally occurring self-organization (Pruessner, 2012; Hesse and Gross, 2014; Cocchi et al., 2017) and there is considerable risk of false positives (Taylor et al., 2013; Hartley et al., 2014; Touboul and Destexhe, 2017; Priesemann and Shriki, 2018), asserting criticality in a *new* system or with a *new* tool requires extraordinary evidence. Since this is a new tool, we created four criteria and set quantifiable standards for concluding a system is consistent with criticality based on avalanche power-laws and we completed this exhaustive battery of tests with shape collapse, a geometrical analysis of self-similarity in the avalanche profiles (see Methods: Experimental Design and Statistical Analysis).

In brief, we found that both the size and duration distributions of the fluctuations treated as avalanches were consistent with power laws (**Figure 2C**), *P*(*A*) ∝ *A*^−*τ*^ and *P*(*D*) ∝ *D*^−*β*^ matching widely reported exponents (Beggs and Plenz, 2003; Priesemann et al., 2009; Hahn et al., 2010; Klaus et al., 2011; Friedman et al., 2012; Shriki et al., 2013; Priesemann et al., 2014; Arviv et al., 2015; Shew et al., 2015; Karimipanah et al., 2017b), obeyed the scaling relation (**Figure 2D**), and exhibited shape collapse, (**Figure 2E**).

Specifically, of the 51 recording groups featuring data from the first 20-minute period of recording from one cell, 98% had power laws in both size and duration distributions. The exponent values for the size distribution were τ = 1.91 ± 0.38 (median ± standard deviation). Exponent values for the duration distribution were *β* = 2.06 ± 0.48. Of the 51 neurons with a recording group from the first 20-minutes, 18 had an additional 20-minute period spanning multiple recordings. All of these 18 groups had power-laws in both size and duration, the exponent values for the size distribution were *τ* = 1.87 ± 0.29 and the exponent values for the duration distribution were *β* = 2.21 ± 0.39.

It is also important to confirm that power-law behavior extends across several orders of magnitude of avalanche durations. We typically demonstrate a power-law distribution over 2.45 ± 0.39 orders of magnitude of duration. For the scaling relation we find a larger span with 2.62 ± 0.23 orders of magnitude across our whole avalanche duration range.

Another statistic crucial to signatures of criticality measures the relationship *between* the power-laws describing size and duration of avalanches (Sethna et al., 2001; Beggs and Timme, 2012; Friedman et al., 2012). If the average avalanche size also scales with duration according to 〈*A*〉(*D*) ∝ *D^γ^*, then the exponent *γ* is not independent, but rather depends on the exponents *τ* and *β* according to *γ* = (*β* − 1)/(*τ* − 1) irrespective of criticality (Scarpetta et al., 2018). For critical systems this condition is enforced because avalanche profiles follows the same shape for all durations which means that this prediction is believed to be more precise than for non-critical systems and the exact values are important (Sethna et al., 2001; Nishimori and Ortiz, 2011). We found that average avalanche size scaled with duration 〈*A*〉(*D*)~*D^γ^* according to a power law and that the observed values of *τ* and *β* provided a good prediction *γ* = (*β* − 1)/(*τ* − 1) of the fitted *γ* (**Figure 2D**).

Specifically, of the 51 recording groups from the first 20-minute period, the fitted scaling relation exponents were *γ_f_* = 1.19 ± 0.05, and the predicted scaling relation exponents were *γ_p_* = 1.17 ± 0.35. For the additional second 20-minute period (18 groups/neurons), the fitted scaling relation exponents were *γ_f_* = 1.21 ± 0.05, and the predicted scaling relation exponents were *γ_p_* = 1.28 ± 0.21.

To affect a more convincing analysis, we defined four stringent criteria that must be independently satisfied before any set of avalanches can be deemed consistent with network dynamics near a critical point (see Methods: Experimental Design and Statistical Analysis). Overall, of the 69 groups of recordings (which includes 18 out of 51 cells twice), 98.6% had power-laws in both the size and duration distributions of avalanches and 92.8% had scaling relations which were well fit by power-laws (*R*^2^ > 0.95). All were deemed non-trivial by the test (*γ_f_* − 1) > *σ_γ_f__* where *σ_γ_f__* is the dataset standard error; *σ_γ_f__* = 0.051. The smallest value was *γ_f_* = 1.094. The fourth constraint, that the *R*^2^ of the predicted scaling relation was within 10% of the best fit scaling relation, was satisfied 85.6% of the time. Together, this set of criteria cannot measure distance from a critical point nor eliminate false positives. However, the take away is that 81% of all recording groups examined were judged to be consistent with network activity near a critical point.

Separating out results: 76% of the 51 recording groups from the first 20-minute period, and 94% of the recording groups from the second 20-minute period were judged consistent with criticality. The general pattern is that the first 20-minute period and the second are both consistent with criticality, but the second group meets our criteria much more frequently. This could be an effect related to the length of time we are able to maintain a patch, or it could be that a better patching results in both longer stable recording ability and better inference of dynamical network properties.

To further discount the possibility of false positives we investigated whether the avalanches within our data set exhibited “shape collapse” (**Figure 2E**). The scaling relation is a consequence of self-similarity (Sethna et al., 2001; Papanikolaou et al., 2011; Friedman et al., 2012; Marshall et al., 2016; Shaukat and Thivierge, 2016; Cocchi et al., 2017). In other words, avalanches all have the same “hump shape” no matter how long they last, this shape is called the scaling-function or avalanche profile. The shape collapse also provides an independent estimate of the scaling relation exponent *γ*, if the estimated exponent, *γ_SC_*, matches the fitted exponent, *γ_f_*, it is considered strong evidence of critical point behavior. For critical systems, the average avalanche profile of an avalanche of duration *D* is given as 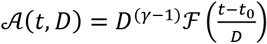. Where *D*^(*γ*−1)^ is a coefficient governing the scaling of height with duration, and 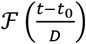 is the scaling-function which describes the universal shape of an avalanche at any duration. The similarity of avalanche profiles of different durations is qualitatively judged (Sethna et al., 2001; Beggs and Plenz, 2003; Friedman et al., 2012; Pruessner, 2012; Timme et al., 2016) by plotting empirically estimated scaling functions for several durations on top of one another after they have been rescaled as part of the process of estimating *γ_SC_*.

We obtained shape collapse across more than one order of magnitude (between about 50 ms to 700 ms) of avalanche durations. Below 50 ms distinct peaks arose and above 700 ms the profile height grew faster than the power-law scaling that worked for shorter duration avalanches. When comparing to plausible alternatives to V_m_ in later sections, we included analysis of mean curvature and avalanche profile peak height along with visual inspection of shape collapse quality (**Figure 2E**). The shape collapse plots begin with short avalanches (20 ms) that are below the median lower cut-off for power-law behavior (which was 256 ms) but are well predicted by the scaling relation.

The exponents estimated from the shape collapse were a good match for both the predicted and fitted scaling relation exponents. The groups of recordings from the first 20 minutes yielded *γ_SC_* = 1.1868 ± 0.042. The average matched absolute percent error was 1.3% with respect to *γ_f_*. A matched signed rank difference of median test revealed that *γ_f_* was not significantly different from *γ_SC_*, simple difference effect size *r_SDF_* = 0.089, p-value *p* = 0.063 (*p* < 0.05 indicates that they are different).

This stage of the analysis showed that, when fluctuations of V_m_ are treated like neuronal avalanches, they are consistent with criticality by the standards of power-laws governing size and duration. We also showed that V_m_ avalanches exhibit geometrical self-similarity across more than one order of magnitude. These factors showed that the cortical circuits driving fluctuations of membrane potential are consistent with a critical branching network according to standards of self-similarity. In our next investigation we compared to population data from microelectrode arrays and other results from literature to test whether V_m_ fluctuations are consistent with the universality requirement of critical branching networks, and also whether they can be used to measure dynamical network properties.

### Membrane Potential Fluctuations are Consistent with Avalanches from Previously Obtained Microelectrode Array LFP Recordings

Importantly, we sought to interpret our results from the analysis of single-neuron V_m_ fluctuations in the context of the more commonly used analysis of multi-unit spiking activity (Friedman et al., 2012; Shew et al., 2015; Marshall et al., 2016; Karimipanah et al., 2017b) or multi-site local field potential (LFP) event detection from microelectrode array (MEA) data (also known as “multielectrode array”) (Beggs and Plenz, 2003; Shew et al., 2015).

In a previous study, avalanche analysis was performed on LFP multi-site MEA recordings from the visual cortex of a different set of 13 ex-vivo eye-attached whole-brain preparations in turtle (Shew et al., 2015). Avalanches were inferred from the steady state (after on response transients but before off response transients) of responses to visual presentation of naturalistic movies as opposed to the resting state activity between presentations (which is where the V_m_ data come from). Avalanche size and duration distributions followed power laws.

The median exponents were *τ* = 1.94 ± 0.27 for the avalanche size distributions and *β* = 2.14 ± 0.32 for the avalanche duration distributions (**Figure 3A**). A scaling relation existed with average exponent *γ_f_* = 1.20 ± 0.06 fitted to the data and *γ_p_* = 1.19 ± 0.07 from the average of the predicted scaling based on theory. The scaling power-law extended over 1-2 orders of magnitude.

**Figure 3:**
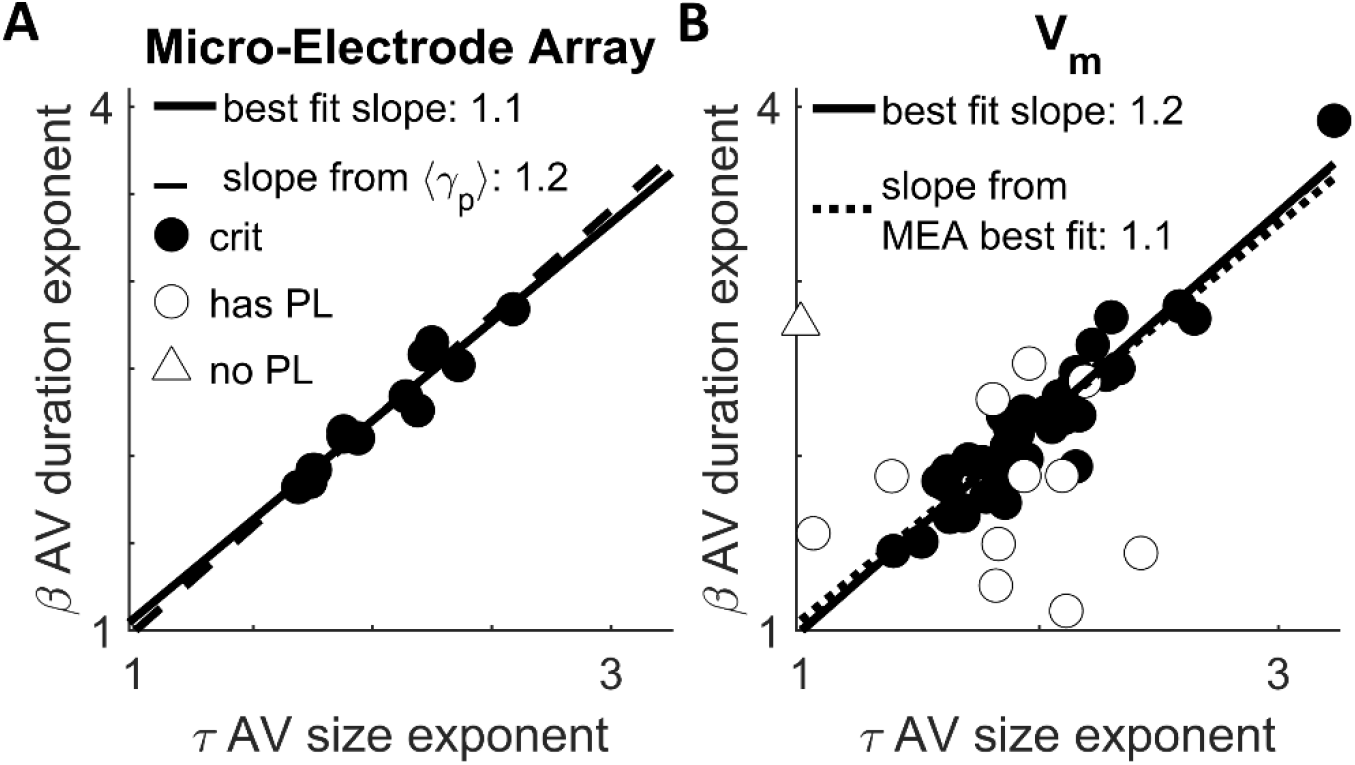
Membrane potential fluctuations are consistent with avalanches from previously obtained microelectrode array data. A plot of the exponents governing power-law scaling of avalanche duration vs the exponents governing avalanche size. Circles indicate data which was best fit to a power-law in both its size and duration. Triangle indicates otherwise (the MLE estimation of a would-be power-law fit, the “scaling index”, is plotted in that case (Jeżewski, 2004)). Filled circles indicate data that meet all four standardized criteria for judging data to be consistent with criticality. **Panel A** is a reproduction from (Shew et al., 2015). It shows the results of avalanche analysis on microelectrode array data collected during the steady state of stimulus presentation in an otherwise identical experimental preparation. The exponent values appear to covary to maintain a stable value of the scaling relation 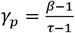. The correlation between *β* and *τ* was high (see Results: The Predicted Scaling Relation Exponent is More Stable than Avalanche Size or Duration Exponents). **Panel B** shows the results of avalanche analysis performed on fluctuations in subthreshold membrane potential. We found power-laws with closely matching exponents and the same scaling relation with the similar level of stability. The correlation between *β* and *τ* was high (see Results: The Predicted Scaling Relation Exponent is More Stable than Avalanche Size or Duration Exponents).

The set of avalanche size, duration, and scaling relation exponents obtained from membrane potential fluctuations (**Figure 3B**) were not distinguishable from the MEA obtained set. The fitted scaling relation exponent *γ_f_* had the least variability of all three kinds of exponents so it is the most likely to show a difference. Thus, if a difference is not significant it suggests universality more strongly than for the avalanche size *τ* or duration *β* distribution exponents.

When we limited our analysis to the first twenty-minute period which contained multiple recordings (51 cells), neither the fitted scaling relation exponent, nor the predicted scaling relation exponent were significantly different from the MEA results. The Wilcoxon rank-sum difference of medians test against the MEA data yielded (*r_SDF_* = 0.164, *p* = 0.37), and (*r_SDF_* = 0.08,*p* = 0.67) respectively. The median exponent values for the size and duration distributions were not significantly different from the median of the MEA data (*r_SDF_* = 0.164, *p* = 0.37) and (*r_SDF_* = 204, *p* = 0.265) respectively.

These results establish V_m_ fluctuations as an informative gauge of high-dimensional information, while also demonstrating that the power-law characteristics are universal properties of the brain, by showing a close match between data at different scales and under different conditions. Further underscoring universality, our results are also similar to the critical exponents measured from other animals such as the *τ* = 1.8 result from in-vivo anesthetized cats (Hahn et al., 2010), though an exhaustive literature search was not conducted, others have conducted incomplete surveys (Ribeiro et al., 2010; Priesemann et al., 2014).

### The Single-Neuron Estimate of Network Dynamics is Optimized at the Network Critical Point

To gain a deeper insight into the relation between single-neuron input and network activity, we investigated a model network of probabilistic integrate and fire model neurons (Kinouchi and Copelli, 2006; Larremore et al., 2011b; Larremore et al., 2011a; Larremore et al., 2012; Larremore et al., 2014; Karimipanah et al., 2017b; Karimipanah et al., 2017a). This model network contains fundamental features of cortical populations, such as low connectivity, inhibition, and spiking, while being sufficiently tractable for mathematical analysis (see Methods: Model Simulations).

In brief, the model network consists of *N* = 10^4^ binary probabilistic model neurons (**Figure 4A**). The connection probability *c* results in a mean in-degree and out-degree of *cN*. The connection strength from neuron *j* to neuron *i* is quantified in terms of the network adjacency matrix *W*. Each connection strength is *W_ij_* drawn from a distribution of (initially) positive numbers with mean η, where the distribution is uniform on [0,2*η*]. A fraction *χ* of the neurons are designated as inhibitory, i.e., their outgoing connections are made negative. The binary state *S_i_*(*t*) of neuron *i* is updated according to 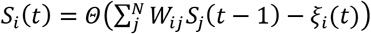, where *ξ_i_*(*t*) is a random number between 0 and 1 drawn from a uniform distribution, and *Θ* is the Heaviside step function.

**Figure 4:**
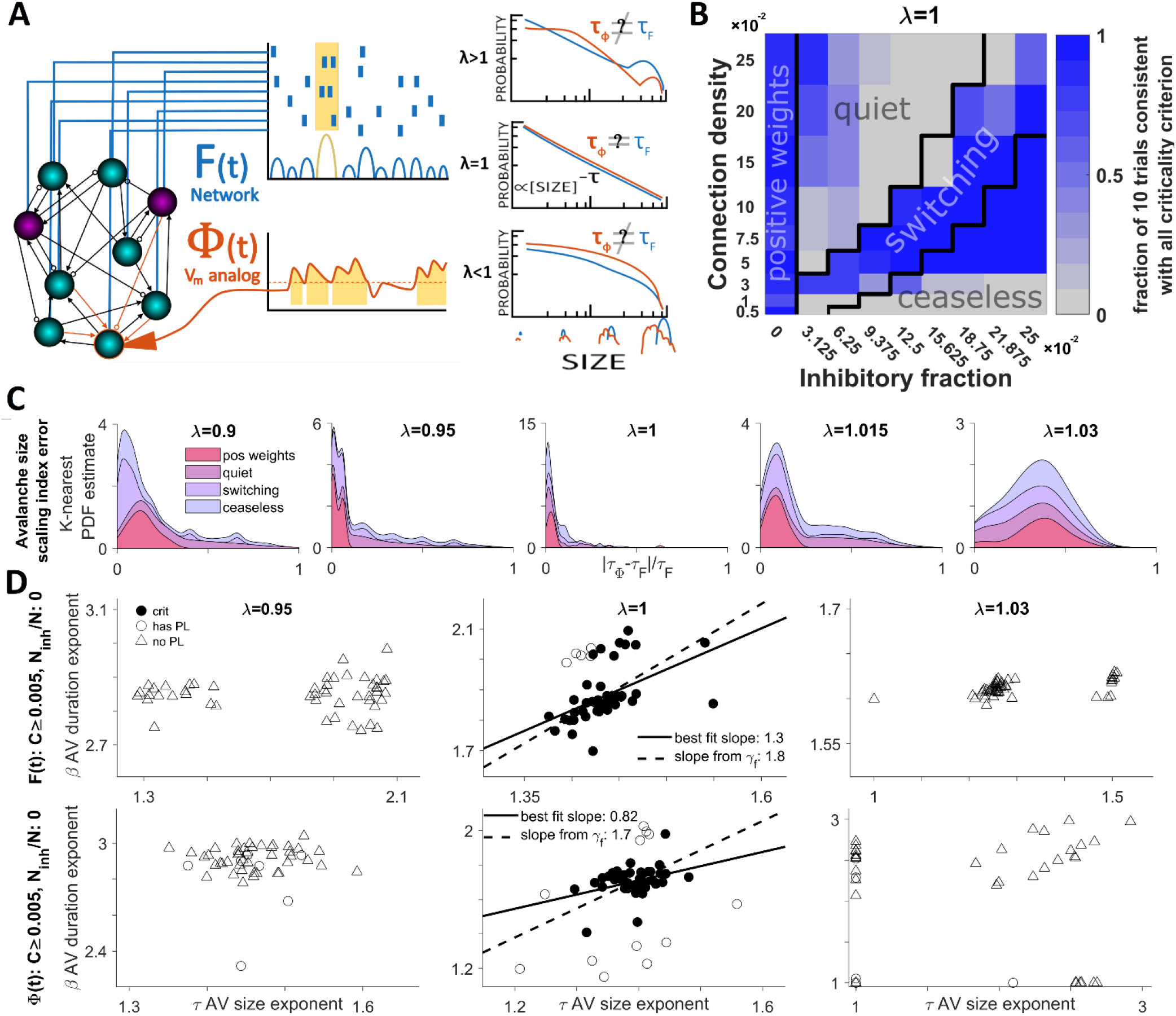
The single-neuron estimate of network dynamics is optimized at the network critical point. **Panel A** illustrates model network consists of 10^4^ excitatory (cyan) and inhibitory (magenta) model neurons with sparse connectivity (line tips: arrows = excitation; circles = inhibition). The simulated model activity (raster plot) is reresented in terms of the single-neuron spiking (raster plot) and the active fraction of the network *F*(*t*) = *S*(*t*)/*N* where population spiking is *S*(*t*). Concurrently, the smoothed inputs (orange) to a single neuron represents the V_m_ proxy, Φ_*i*_(*t*). The threshold (dashed line) crossings of Φ_*i*_(*t*) define avalanches (see Methods: Experimental Design and Statistical Analysis). Avalanches of *F*(*t*) and Φ_*i*_(*t*) are analyzed in terms of their size (shown) and duration (not shown) distributions and their corresponding exponents, *τ*. Avalanche statistics depend on several network parameters including the critical branching tuning parameter *λ*. **Panel B** shows how the inclusion of inhibition affects the network behavior. The black lines mark the boundaries of arbitrarily defined parameter regions roughly corresponding to distinct kinds of behavior. The shade of blue indicates what fraction of ten trials at each point met all four of our standardized criteria for consistency with expectations of critical branching behavior. **Panel C** is a stacked area chart showing the probability density distribution of size exponent error (between *F*(*t*) and Φ_*i*_(*t*)) for different *λ* and dynamical regimes. The vertical thickness of each color band shows the probability density for that subset of the data while the outer envelope shows the over-all probability density. Probability density is estimated with a normal kernel smoothing function. In this panel we can see that power-law scaling is most similar at criticality despite variability dependent on the parameter regime. **Panel D** shows a complete summary of the tests for criticality when applied to *F*(*t*) (top row) and Φ_*i*_(*t*) (bottom row). From this we can confirm that the system is consistent with criticality when there is no inhibition. The subsampling method Φ_*i*_(*t*) demonstrates consistency with criticality but displays a wider dispersion of exponent estimates. For experimental V_m_ and MEA data there was a large correlation between *β* and *τ* showing that the scaling relation (which predicts the slope of the trendline) is much more stable than exponent values. This is not the case for the model where for *F*(*t*) the correlation is low (see Results: The Predicted Scaling Relation Exponent is More Stable than Avalanche Size or Duration Exponents).

The largest eigenvalue *λ* = *ηcN*(1 − 2*χ*) of the network adjacency matrix *W*, characterizes the network dynamics, with critical network dynamics occurring at *λ* = 1. This tuning parameter *λ* controls the degree to which spike propagation “branches”: *λ* = 1 means that one spike creates one other spike on average, *λ* > 1 implies that one spike creates more than one other spike while *λ* < 1 means that one spike creates less than one other spike (Haldeman and Beggs, 2005; Kinouchi and Copelli, 2006; Levina et al., 2007; Larremore et al., 2011b; Larremore et al., 2012; Kello, 2013; Larremore et al., 2014).

The input to model neuron *i*, is 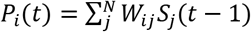 and provides the link between network activity and single-neuron activity. From this we can derive a simple mathematical result characterizing how estimation of network properties is optimized at criticality.

If we let *K_i_*(*t* − 1) denote the number of active neurons *in the presynaptic population* of neuron i, then we can rewrite the input to a model neuron as a sum of independent and identically distributed random variables drawn from the non-zero entries of 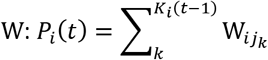. After implementing inhibition by inverting some elements of *W* the distribution of weights is not uniform but piecewise uniform. Weights are drawn uniformly from the interval [−2*η*, 0] with probability *χ* and from the interval [0,2*η*] with probability 1 − *χ*. The mean of the nonzero entries of W are denoted with a prime so that the mean is 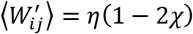 and the standard deviation is 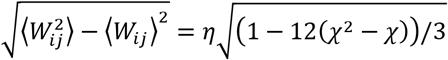. Now we can find the mean behavior of the input integration function as it relates to the presynaptic population:

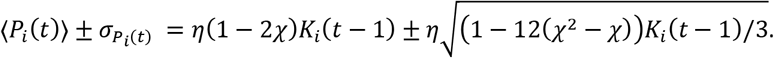

We learn three things by examining the mean behavior of the input integration function. First, the mean grows as *O*(*K_i_*) but the standard deviation grows as the root 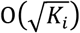, so the function becomes a more precise estimator of network activity with increasing activity in the presynaptic population (increasing *K_i_*). Second, the input integration function *P_i_*(*t*), is rarely negative. At the parameter combination *c* = 0.005 and *χ* = 0.25 (which has the largest variance relative to the mean) the mean becomes more than one standard deviation larger than zero when *K_i_* > 5. Third, and most importantly, the input integration function is an averaging operator and the tuning parameter *λ* biases that averaging operation. To show this we only need two observations: the instantaneous firing rate averaged over the presynaptic population is the number of active neurons divided by the expected total number of presynaptic neurons, *ω_i_*(*t*) = *K_i_*(*t*)/*cN*. Next, we rearrange the definition of lambda to get *λ*/*cN* = *η*(1 − 2*λ*). Substituting these two observations into the mean behavior of our input integration function we get the key result:

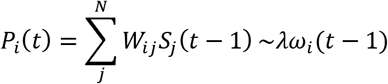

Note that 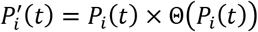 directly represents the probability for the neuron to spike at time *t*.

These results demonstrate that the inputs to a neuron *P_i_*, and the instantaneous firing rate of that neuron are the result of an averaging operator acting on the presynaptic population, which is a subsample of the network. Furthermore, the tuning parameter *λ* not only modulates the relationship of single neuron firing to downstream events (also known as branching), but also governs how the input to a neuron relates to the presynaptic population. It biases the averaging operator to either amplify firing rate (*λ* > 1) or dampen it (*λ* < 1). Therefore, our model implements both critical branching and the inverse of the critical branching condition, a *critical coarse-graining* condition. The model is a network of subsampling operators who only capture whole-system statistics when *λ* = 1 and the operators reflect an unbiased stochastic estimate of mean firing rate among the subsample (the presynaptic population).

To further evaluate the relation between single-neuron input and network activity under different conditions, we simulated the described network of 10^4^ model neurons for a total of 405 different parameter combinations, including connection probability, inhibition, and maximum eigenvalue (**Figure 4A**), each parameter combination was repeated ten times. We then compared the avalanche analysis results of simulated network activity 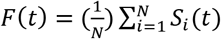 and the input to a single neuron (the input integration function). However, 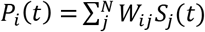 is the probability that neuron *i* will fire at time *t*, also known as the instantaneous firing rate of neuron *i*.

Membrane potential is not a direct representation of firing rate, but rather the firing rate is related to synaptic input through the F-I curve which is non-linearly related to membrane potential. This non-linearity could destroy the correspondence between the simulated single neuron signal and network activity. In order to better facilitate comparison of the simulated input integration function with the experimentally recorded membrane potential, we constructed a proxy for the subthreshold membrane potential, Φ_*i*_(*t*), of a model neuron by convolving the simulated input *P_i_*(*t*) with an alpha function (see Methods: Model Simulations).

The parameter space has four distinct patterns of critical network behavior (**Figure 4B**). Qualitatively, these were reflected in the network activity. As the presence of these paradoxical behaviors may indicate the presence of second phase-transition tuned by the balance of excitation to inhibition (Shew et al., 2011; Poil et al., 2012; Kello, 2013; Hesse and Gross, 2014; Larremore et al., 2014; Scarpetta et al., 2018) several key results differ strongly and thus are reported separately for these regions of parameter space.

These regions are defined in terms of the connection density and inhibition and shown in **figure 4B**. First is the “positive weights” region, there is no inhibition (*χ* = 0) and the network is a standard critical branching network. The second region, “quiet”, has a small increase in the fraction of inhibitory neurons. Activity lasts slightly longer than for the classically critical network. The third region is called the “switching” regime because network activity switches between a low mean and a high mean (like “up and down states” (Destexhe et al., 2003; Millman et al., 2010; Larremore et al., 2014; Scarpetta et al., 2018)). This occurred in the middle portion of the values of connectivity and inhibition. Lastly, we have the “ceaseless” region, with a large fraction of inhibition, relative to connection density, activity never dies out. This region is defined by *c* < (10*e*^12*χ*^ − 13)/100 and *χ* > 0. Three of these regimes are displayed in figure 5A, the “quiet” regime” is mostly redundant to the “positive weights” region.

**Figure 5:**
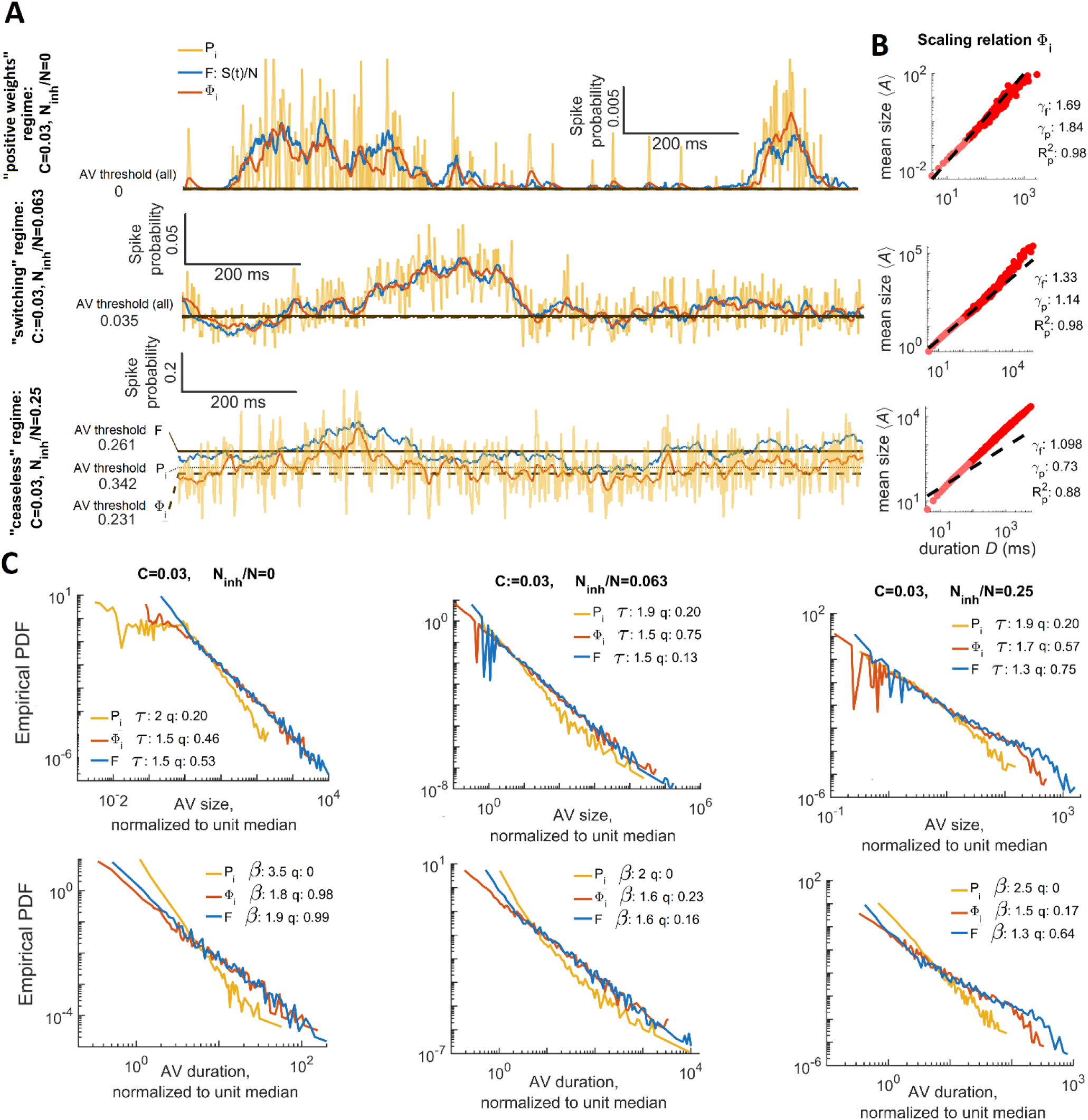
Inputs to a neuron stochastically estimate firing of its presynaptic pool in this critical branching model. **Panel A** shows differences in model activity dynamics with parameter regions (constant connectivity, *λ* = 1, but inhibition, *χ* varies). Each plot shows the active fraction of the network *F*(*t*) in blue, the instantaneous firing rate of node, *P_i_*(*t*), is in gold and the V_m_ proxy for the same node, Φ_*i*_(*t*), is in orange. The node is randomly selected from the nodes with degree within 10% of mean degree. The V_m_ proxy is produced by convolving the firing rate of a single neuron with an alpha function with a 4 ms time constant. The top plot shows that with no inhibition (or very little inhibition) activity in this parameter region dies away to zero and is unimodally distributed about a small value. The middle plot shows that moderate amounts of inhibition results in self-sustained activity that is bimodally distributed about one high and one low value. The bottom plot shows that when the fraction of nodes that are inhibitory is much larger than connection density activity is self-sustaining and unimodally distributed about a high value with low variance relative to the mean. **Panel B** shows the scaling relation for the avalanches inferred from Φ_*i*_(*t*) at different levels of inhibition, as in panel A. Inhibition detrimentally impacts the validity of the scaling relation predictions, which are required for consistency with critical branching. The predicted (*γ_p_*) and fitted (*γ_p_*) scaling exponents are indicated as is the goodness of fit (Æp) for the predicted exponent. **Panel C** shows how avalanche (fluctuation) statistics vary with the parameter set displayed in panels A and B. The top row shows avalanche (fluctuation) sizes, while the bottom row shows the duration distributions. Exponents *τ* (size distribution) and *β* (duration distribution) as well as comparison quotients *q* are annotated on the plot. From these plots, we can see that temporal smoothing (Φ_*i*_(*t*)) is necessary to accurately capture *F*(*t*). Additionally, we see that mismatch between the *F*(*t*) and *P_i_*(*t*) avalanche distributions vary with network parameters. At high levels of inhibition, the *i*(*t*) avalanches are power-law distributed over smaller portions of their support. For Φ_*i*_(*t*), neither of the networks with less inhibition show the cutoffs associated with under sampling a critical branching network.

We looked at the magnitude of relative error between estimated exponents for the avalanche size distribution (**Figure 4C**) to determine how well our proxy neural inputs, *ϕ_i_*(*t*), reflected network activity, *F*(*t*), in different parameter regions, and with different values for the tuning parameter, *λ*. Importantly the least error occurred for *λ* = 1 with and without the presence of inhibitory nodes. This insensitivity to parameter differences supports the claim (Larremore et al., 2014) that the system becomes critical when *λ* = 1 even in the presence of inhibition.

However, the four regions of parameter space perform differently according to our four standardized criteria for consistency with criticality. In the “positive weights” region 90% of 90 trials (nine points in parameter space with ten trials per point) have network activity that meets all four criteria when the tuning parameter is set at criticality (*λ* = 1) (**Figure 4C**). Meanwhile 39% meet the criteria in the “ceaseless” region, 19% do in the “quiet” region, and 67% do in the “switching” region which may indicate the location of a second phase-transition and shows that evidence for precise criticality in this model is limited once inhibition is included.

As we vary the tuning parameter, we can clearly distinguish critical from non-critical systems. Over all 47% percent of trials meet all four criteria when *λ* = 1, while 3% do when *λ* = 0.95, 18% do when *λ* = 1.015, 1% do when *λ* = 0.9, and 1% do when *λ* = 1.03 (**Figure 4D**).

The estimated power-law exponents show that the avalanche size distributions for *F*(*t*), *P_i_*(*t*), and Φ_*i*_(*t*) are most alike at criticality. Note that estimated exponents serves as the “scaling index”, a measure of the heavy tail even when a power-law is not the statistical model that fits best (Jeżewski, 2004). The fact that matching between network activity and the input integration function was best at criticality is important because it underscores the scale-free nature of critical phenomena and contrasts with the results obtained when testing a different relationship between subsampling methods and network structure (Priesemann et al., 2009; Yu et al., 2014; Levina and Priesemann, 2017).

While the system was both critical (*λ* = 1) and in the positive weights region, our V_m_ proxy Φ_*i*_(*t*) met all four criteria for consistency with criticality 74% of the time for 90 trials (**Figure 4D**) while *P_i_*(*t*) met all four only 1% of the time. The network activity had avalanche size and duration exponent values *τ_F_* = 1.43 ± 0.04, and *β_F_* = 1.87 ± 0.09, (Figure 4D) and had a fitted scaling relation exponent, *γ_F_f__* = 1.83 ± 0.02, and a predicted exponent *γ_F_p__* = 1.99 ± 0.23. The membrane potential proxy, Φ_*i*_(*t*) had slightly lower avalanche size and duration exponent values that fluctuated around the paired network values, *τ*_Φ_ = 1.40 ± 0.06, and *β*_Φ_ = 1.73 ± 0.17, (Figure 4D) and exclusively lower scaling relation exponents *γ*_Φ_*f*__ = 1.68 ± 0.02. While the unsmoothed *P_i_*(*t*) varied considerably more it had size and duration exponents that were almost exclusively higher than the paired network values, *τ*_p_ = 1.87 ± 0.50, and *β*_P_ = 2.84 ± 1.45, with a fitted scaling relation exponent that was exclusively lower *γ*_P_*f*__ = 1.68 ± 0.02.

In figure 5, we compared different population dynamics estimation techniques by looking at avalanches inferred from *P_i_*(*t*) (the inputs to neuron *i*), and the V_m_ proxy Φ_*i*_(*t*). Both *P_i_*(*t*) and Φ_*i*_(*t*) fluctuate about *F*(*t*) but *P_i_*(*t*) is much noisier (**Figure 5A**), in the ceaseless regime *P_i_*(*t*) and Φ_*i*_(*t*) are systematically offset. Avalanches inferred from Φ_*i*_(*t*) had average sizes that scaled with duration (Figure 5B). Avalanches from Φ_*i*_(*t*) consistently had duration and size distribution exponents that were closer to network avalanches than avalanches from *P_i_*(*t*). However, *P_i_*(*t*) performed satisfactorily in the sense that its error was systematically offset and best at criticality (**Figure 5C**).

Including inhibition introduces several important differences. For the ceaseless region with *λ* = 1, far fewer trails meet our criteria, however *P_i_*(*t*) follows *F*(*t*) much more closely. The network activity had avalanche size and duration exponent values *τ_f_* = 1.48 ± 0.09, and *β_F_* = 1.53 ± 0.09, and had a fitted scaling relation exponent, *γ_F_f__* = 1.23 ± 0.11. The membrane potential proxy, Φ_*i*_(*t*) had slightly higher avalanche size and duration exponent values that fluctuated around the paired network values, *τ*_Φ_ = 1.51 ± 0.19, and *β*_Φ_ = 1.57 ± 0.17, but nearly identical scaling relation exponents *γ*_Φ_ = 1.23 ± 0.11. While the unsmoothed *P_i_*(*t*) varied considerably more, it had size and duration exponents that were almost exclusively higher than the paired network values, *τ*_p_ = 1.88 ± 0.20, and *β*_P_ = 2.18 ± 0.34, with a fitted scaling relation exponent that was slightly lower *γ*_P_*f*__ = 1.19 ± 0.07.

When *γ* ≠ 1 both Φ_*i*_(*t*), and *P_i_*(*t*) failed to meet all four criteria for criticality at the same high rate as **F*(*t*)* (to within 1%). This lack of false positives confirms that these signals are useful for characterizing critical branching. In figure 4, panel B, we calculated the absolute magnitude of relative error between the size exponent from avalanche analysis performed on *F*(*t*) and Φ_*i*_(*t*). As expected, the avalanches were usually not power-laws according to our standards, in this case the exponent is known as the “scaling index” and describes the decay of the distribution’s heavy tail (Jeżewski, 2004).

When we set *λ* = 0.95 we see a moderate deterioration in the ability of either Φ_*i*_(*t*) or *P_i_*(*t*) to recapitulate network exponent values. The error is no longer systematic; thus, they cannot be used to predict network values. The variability of the exponents increases greatly for Φ_*i*_(*t*) while it decreases for *P_i_*(*t*). The exponent error increases slightly over the *λ* = 1 and the base of the distribution is much broader.

Reducing *λ* further, to *λ* = 0.90, the input integration function, *P_i_*(*t*)~*λω_i_*(*t* − 1), rapidly dampens impulses (wi is the instantaneous firing rate over the presynaptic population for neuron *i*). Variability continues to increase, and a systematic offset does not return. Exponent error is now much broader. With branching this low, events often are not able to propagate to the randomly selected neuron, an exception is the “ceaseless” regime where activity is still long lived.

When we set *λ* = 1.015 we see a dramatic deterioration in the ability of either Φ_*i*_(*t*) or *P_i_*(*t*) to recapitulate network values. Variability in exponent estimation increases for both Φ_*i*_(*t*) and *P_i_*(*t*). Exponent error increases rapidly, underscoring the inability to estimate network activity from neuron inputs.

Increasing *λ* further to *λ* = 1.03 produces an input integration function, *P_i_*(*t*)~*λω_i_*(*t* − 1), that rapidly amplifies all impulses and the network saturates. The effect is that variability in the estimated exponents decreases and a systematic offset returns, with both Φ_*i*_(*t*) and *P_i_*(*t*) producing exponents that are exclusively and considerably higher than network values. Exponent error reveals that estimating network properties from the inputs to a neuron is probably not possible for supercriticality in this model.

The results here show that the V_m_ proxy represents an effective way of subsampling network flow. This is a hallmark of the near-critical region in the PIF model and a manifestation of scale-freeness. Criticality in our model corresponds to the point when the inputs to a neuron represent an average of the activity of the presynaptic population. Importantly we explored why it works, as well as showing that it does work in experimental data. This analysis, presented in forthcoming sections, uncovered that proper temporal and spatial aggregation is important as is the role of inhibition in membrane potential dynamics. This supports both the criticality hypothesis, and tight balance (Boerlin et al., 2013; Barrett et al., 2014; Denève and Machens, 2016). Additionally, it has specific implications for the information content of membrane potential.

### The Predicted Scaling Relation Exponent is More Stable than Avalanche Size or Duration Exponents

A key part of the study of criticality in neural systems is the assumption that biological systems must self-organize to a critical point. The precise critical point is a very small target for a self-organizing mechanism in any natural system. So, a key question is whether the self-organizing mechanism of the brain prioritizes efficiently achieving information processing advantages of scale-free covariance at the expense of being slightly sub or super-critical (which is a larger target) (Priesemann et al., 2014; Tomen et al., 2014; Williams-Garcia et al., 2014; Gautam et al., 2015; Clawson et al., 2017).

Our data offered unexpected insight. It is known that so long as three requirements are met the scaling relation will be marginally obeyed: Avalanche size and durations must be power-law distributed (with exponents *τ* and *β* respectively) and average size must scale with duration according to a power-law with exponent *γ*. Given those three requirements one can derive a prediction for the scaling exponent, *γ_p_* = (*β* − 1)/(*τ* − 1) without needing to assume criticality (Scarpetta et al., 2018). However, without any other assumptions one expects *β* and *τ* to be independent so plotting one against the other should make a point-cloud that is symmetrical, not stretched along a trendline (**Figure 3**).

We analyzed the independence of *τ, β*, and *γ* measured from experimental data (where self-organization is hypothesized) and compared it to model data (where self-organization is impossible, but criticality is guaranteed). We found that *β* and *τ* are more independent and the predicted scaling relation is more variable for the model than for experimental data in which *β* and *τ*covary, apparently in order to maintain a fixed scaling relation prediction.

The previous multi-site LFP recordings displayed a range of values for the avalanche size *τ*and duration *β* distribution exponents across the tested brain preparations. Interestingly, the exponent values were not independent, rather the duration exponent varied linearly with the size exponent (Shew et al., 2015) (Figure 3A). The single-neuron V_m_ fluctuations, reported here, produced a similar linear relationship between size and duration exponent (Figure 3B). Algebraic manipulation of the predicted scaling exponent *γ_p_* = (*β* − 1)/(*τ* − 1) provides a clue. If the scaling relation (*β* − 1) = *γ*(*τ* − 1) is obeyed and if is a *fixed universal property*, then the linear relationship *β_j_*~*γ_p_τ_j_* holds across different cells and animals.

To demonstrate this important result, variability in the predicted scaling-relation is much less than expected, we propagate errors and assume independent *β* and *τ*. We would expect the standard deviation of *γ_p_* to be 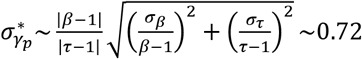 which is roughly twice the real value in V_m_ data, *σ_γ_p__* ~0.35.

Pearson correlation, *ρ*, confirms strong dependence between *τ* and *β, ρ_τβ_* = 0.61, p-value *p* = 2.57 × 10^−6^ for the V_m_ data while for the MEA data *ρ_τβ_* = 0.96, p-value *p* = 1.01 × 10^−7^. From this we confirm what figure 3 shows: the variability in *τ*and *β* are not independent and this implies the existence of an organizing principle connecting *τ* to *β*. Whatever the principle may turn out to be, one of its effects is the maintenance of low variability in *γ_p_* at the expense of greater variability in *τ* and *β*.

A principle reason to suspect self-organization is that this trend is not seen in the model results. Importantly, *τ*and *β* are independent of the scaling-relation exponent function, though still weakly correlated. In this model there is no adaptive organizing principle driving this network to criticality, instead the structure is fixed and set to be at the critical point. This shows how systems behave in the absence of self-organization. No parameter is being maintained at low variability at the expense of other parameters.

Limiting ourselves to simulated network activity for the *λ* = 1 case without inhibition (Figure 4C), propagation of errors leads us to expect the standard deviation of the scaling-relation prediction to be 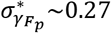 which is very close to real value *σ_γ_F_p___* ~0.23. The correlation is statistically significant at the 5% level, but much smaller *ρ_τβ_* = 0.23, p-value *p* = 0.027.

In conclusion, the linear trend between avalanche size and duration exponents is not a universal property of critical systems because it was not found in the model. This suggests that the linear trend is enforced by an organizing principle at work in the brain but absent in the model. This principle prioritizes maintaining stability in either the scaling of avalanche size with duration, or the power-law scaling of autocorrelation which is closely related to the scaling relation and scale-free covariance via the power-law governing auto-correlation (Bak et al., 1987; Sethna et al., 2001).

### Non-Linearity and Temporal Characteristics such as High-Order Correlation, Proper Combination of Synaptic Events, and Signal Time-Scale are Required to Reproduce Network Measures from Single-electrode Recordings

In order to demonstrate that subthreshold membrane potential fluctuations can be used as an informative gauge of cortical population activity it is necessary to compare against alternative signals which have either been used by experimentalists as a measure of population activity or that share some key features of membrane potential but are missing others. By making these comparisons we can illuminate which features of the membrane potential signal are responsible for its ability to preserve properties of cortical network activity. Additionally, it is necessary to check whether the statistical properties of avalanches can be explained by random processes unrelated to criticality. To address these points of the investigation, we analyzed five surrogate signals: single-site LFP recorded concurrently with the V_m_ recordings, two phase-shuffled versions of V_m_ recordings, computationally inferred excitatory current, and the same inferred excitatory current further transformed to match V_m_ autocorrelation (which tests the role of IPSPs by removing them).

### Negative fluctuations of LFP disagree with V_m_ and MEA results and are inconsistent with critical branching avalanches

The first alternative hypothesis to test is whether the LFP could yield the same results. We used low-pass filtered and inverted single site local field potential (LFP) which is commonly believed to measure local population activity. However, in our analysis it did not recapitulate the results from either MEA or V_m_ avalanche analysis. We obtained viable single-site LFP recordings (see Methods: Extracellular Recordings), simultaneous and adjacent with whole-cell recordings, for 38 of the 51 neurons reported above. We performed avalanche analyses on the LFP recordings using a procedure like the one described for the V_m_ recordings (see Methods: Intracellular Recordings) (**Figure 6**). LFP recordings were grouped the same way V_m_ recordings were in order to match them for comparison. However, the numbers of recordings are not the same because there were two or three cells being patched alongside (within 300 *μm*) one extracellular electrode and there was not always a simultaneous LFP recording. LFP also produced more avalanches per 2-5-minute recording *N_AV_* = 1128 ± 348. The are 23 20-minute periods spanning multiple LFP recordings. These recordings were gathered into groups and matched against 49 V_m_ recording groups (38 from the first 20-minute period, 11 from the second). Additionally, there were 16 20-minute periods spanning only one LFP recording but with more than 500 avalanches. The concurrent V_m_ recordings did not have enough avalanches. This gives us 39 LFP avalanche data sets.

**Figure 6:**
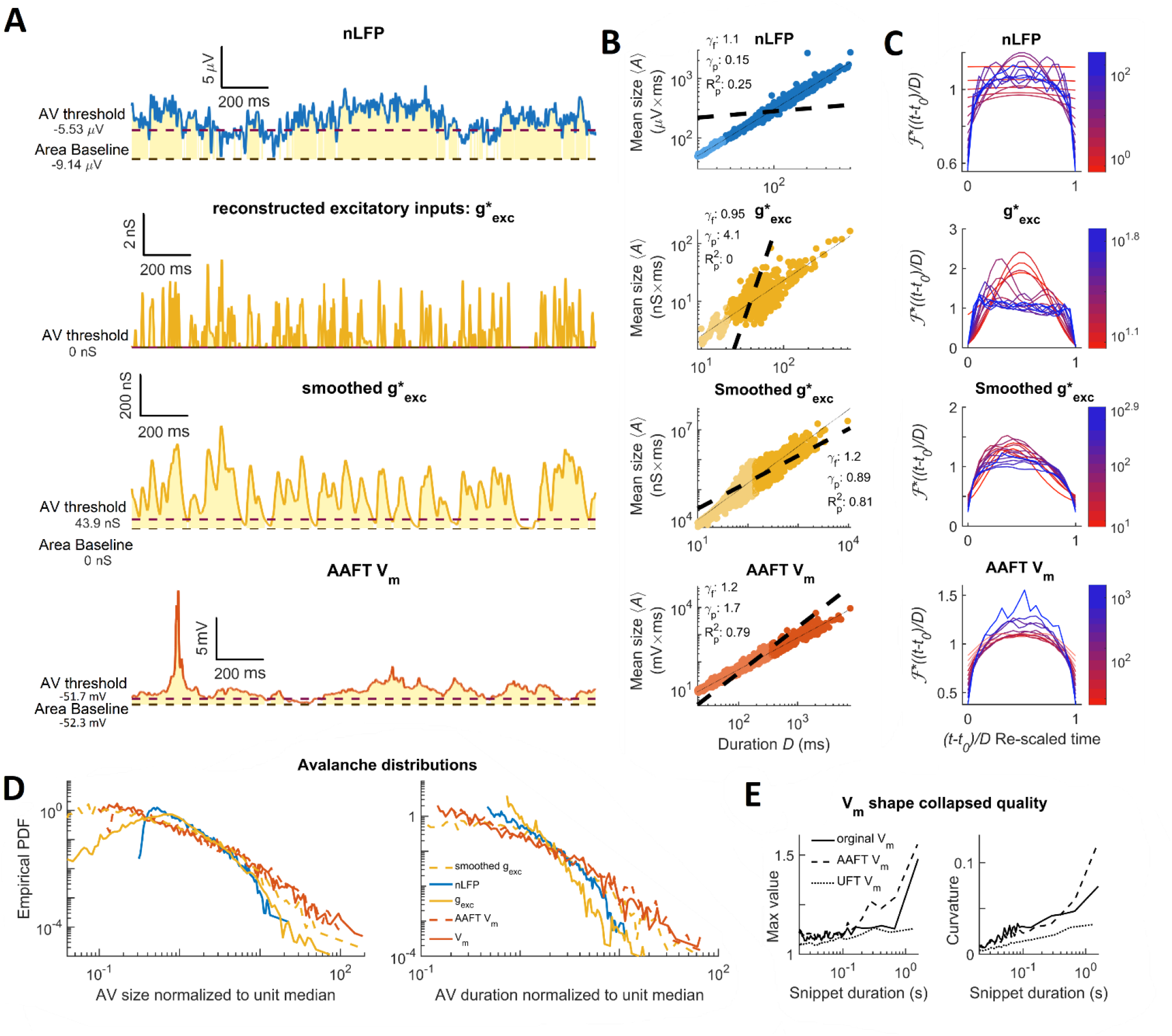
Comparison to surrogate signals reveals the importance of non-linearity and temporal characteristics such as high-order correlation, proper combination of synaptic events, and signal time-scale. **Panel A** shows alternative signals and surrogate data time synchronized to figure 2B and showing thresholds and integration baselines (dashed lines) with avalanche areas marked in yellow. The top row shows the inverted LFP signal. The LFP is low-pass filtered (0-100 Hz), inverted, detrended and analyzed for avalanches identically to membrane potentials. The second and third rows show the inferred excitatory inputs to a neuron. An algorithm reconstructs the timing and shape of ePSPs from V_m_. The resultant signal, 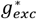, is much faster, making it analogous to the *P_i_*(*t*) signal from the PIF model. This signal is smoothed (third row, see Methods: Model Simulations for details) to produce a signal that is like V_m_ (Figure 2B) would be if it lacked IPSPs. The last row provides an example of amplitude matched phase shuffled surrogate data (amplitude adjusted Fourier transform algorithm). **Panel B** shows the scaling relation in the same order and dataset as panel A. The dashed line is the predicted scaling relation exponent inferred from power-law fits to the size and duration distributions of positive fluctuations. In cases where a power-law is not the best model the exponent nonetheless gives the average slope of a linear regression on a log-log plot, a “scaling index” (Jeżewski, 2004). The predicted (*γ_p_*) and fitted (*γ_f_*) scaling exponents are indicated as is the goodness of fit 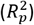 for the predicted exponent. Mean size scales with duration for all signals but often it is trivial (*γ_f_*~1) or poorly explained by a power-law 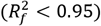, and it is rarely a good match with the prediction from the scaling relation. **Panel C** shows shape collapse from the total dataset in the same order and dataset as panel A. The color indicates the duration according to the scale bar. If self-similarity is present each avalanche profile will collapse onto the same curve: 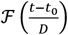. The LFP illustrates a trivial scaling relation that is not produced by true self-similarity: limited curvature and the exponents are very close to one. The second row shows the reconstructed excitatory inputs, 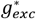, and lacks shape collapse as expected from the lack of a scaling relation power-law in panel B. The third row shows that sensible curvature re-emerges with smoothing but does not produce a universal scaling function. In the last row the phase shuffled V_m_ shows a shape collapse which is worse than for the original V_m_ (Figure 2E). **Panel D** shows size and duration distributions from each signal compared with the V_m_ (in solid red). The phase shuffled V_m_ (dashed red) still obeys power-laws but the exponent values disagree, and it less frequently meets our standardized criteria. Unsmoothed 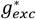(solid gold) is more like inverted LFP than anything else. When 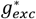 is smoothed (dashed gold) it becomes closer to the original V_m_ but retains pronounced curvature in the duration distribution. We see V_m_, AAFT, and smoothed 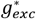 produce distributions which extend over similar orders of magnitude (~2). **Panel E** shows maximum value and curvature of the average profiles after “collapse” as functions of duration. Shape collapse quality is a subjective measure, but these give a more quantitative perspective. Good shape collapse should have a fixed maximum value and a high but fixed mean curvature. For comparison, the UFT (Unwindowed or Unadjusted Fourier Transform) phase shuffled data is also shown to provide a comparison to low curvature but a fixed maximum value. By visual inspection of AAFT and V_m_ it is apparent that the asymmetry is gone and that deviation from the collapsed shape begins at shorter durations. The max value diverges from a linear trend sooner for AAFT (~0.15 seconds, 0.5) than for V_m_ (~0.7 seconds). Curvature also diverges sooner for the AAFT (0.5 seconds vs 0.7 seconds). Curvature does not become appreciable until about 50-70 ms. Between the onset of curvature and divergence of max value there are 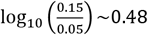 orders of magnitude for AAFT and 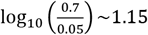 orders of magnitude for the original V_m_.

The LFP recording groups performed poorly according to our four criteria for consistency with criticality. Of the 39 LFP recording groups, only 41% percent had acceptable scaling relation predictions and only 36% met all four standard criteria for criticality (**Figure 7A**). The additional criterion of shape collapse was not observed (**Figure 6C**), there was no linear trend among the exponents governed by the scaling relation and the exponents did not match MEA data (**Figure 3A**). However, 85% produced power-law fits for size and duration, 92% had scaling relations well fit by power-laws and all were non-trivial. We expect from (Touboul and Destexhe, 2017) that some fraction of non-critical data will pass the four standard criteria by chance, so long as the data have a 1/*f* power spectrum.

**Figure 7:**
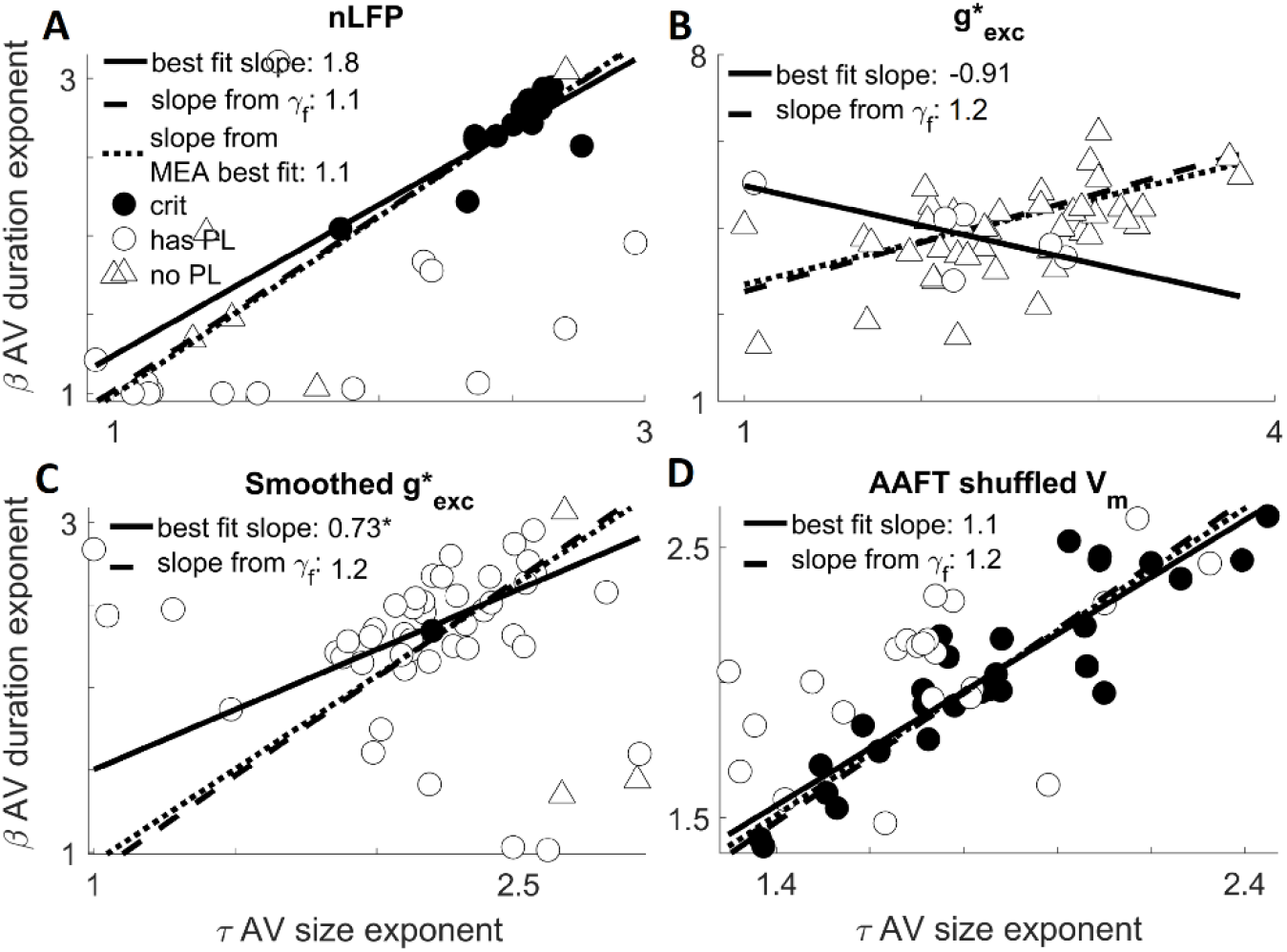
Plausible alternative signals fail to demonstrate consistency with criticality. A plot of the exponents governing power-law scaling of avalanche duration vs the exponents governing avalanche size. Circles indicate data which was best fit to a power-law in both its size and duration. Triangle indicates otherwise (the MLE estimation of a would-be power-law fit, the “scaling index”, is plotted in that case (Jeżewski, 2004)). Filled circles indicate data that meet all four standardized criteria for judging data to be consistent with criticality. We show the performance summary for the first group of data from each cell (the first 20-minute period which contained multiple recordings). The best fit slope is from linear regression to the plotted or indicated data, this is compared to the slope predicted by the mean *γ_f_* (the exponent describing how avalanche size scales with duration). **Panel A** shows that positive fluctuations of inverted LFP were less likely to be power-law distributed and the power-law exponents tended to be unstable and not resemble MEA results. All 39 LFP datasets are represented. **Panel B** shows results from the reconstruction of excitatory input conductance *g_exc_*. Only 12% were power-law distributed. The results do not resemble the MEA results. The slope from the trendline matches the scaling relation exponent but the regression is bad, *R*^2^ = 0.51. **Panel C** shows how adding back some temporal smoothing to 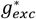 can improve results, 94% have power-laws but the exponents are more variable and generally larger. Most (96%) fail to have scaling relations which are well described by power-laws. The exponents *β* and *τ* are less independent but are not well described by the regression trendlines (*R*^2^ = 0.35). The fit is applied only to the upper right cluster, excluding the outliers in the region *β* < 1.6 and *τ* < 1.6. **Panel D** shows the summary of results from the AAFT phase shuffled V_m_. As expected for a shuffling that preserves autocorrelation, power-laws are also preserved. However, the exponents are shifted down (especially the size distribution exponent). Far more fail to meet our criteria for consistency with criticality, as statistically significant difference (see Results: Stochastic Surrogates are Distinguishable from V_m_ or MEA Results, Reveals Importance of Non-Linear Filtering). Significantly fewer data sets have scaling relations well described by a power-law (75% as opposed to 90%), this is consistent with a slightly worse shape collapse (Figure 6C).

To emphasize that these results are chance we can limit ourselves to just those with the best chance of meeting the scaling relation criteria by picking those that have power-laws in the size and duration distributions. This is enough to expect the scaling relation to be obeyed if mean size scales geometrically with duration (Scarpetta et al., 2018). It is still the case that only 42% of recording groups meet the three remaining standard criteria for consistency with criticality. Therefore, having power-laws is statistically independent of meeting the other criterion for consistency with criticality.

Not only does the single-site LFP data differ from MEA and V_m_ data because it fails to demonstrate consistency with criticality, it is also the case that the scale-free properties which do exist are not representative of the MEA data or the simultaneous V_m_ recordings. The failure was not because LFP recordings co-occurred with decreased consistency with criticality more generally. Eighty-one percent of the matched V_m_ recordings met all the criteria, while 58% of the LFP recordings did, a statistically significant dissimilarity, odds ratio (*r_OR_* = 7.65 with *p* = 1.1 × 10^−5^).

The estimated exponents from all 39 LFP recording groups were highly variable. The duration distribution and scaling relation were most dissimilar to V_m_ and MEA data. Of the 33 LFP groups which were power-law distributed, the avalanche size exponent had a median value *τ* = 1.90 ± 0.63 while the duration exponent was *β* = 1.41 ± 0.9 (very low) (**Figure 7A**) and the fitted exponent was *Yf = 1.11* ± *0.02*. The predicted scaling-relation exponents were inaccurate with *γ_p_* = 0.89 ± 0.76 for the subset of recording groups which had power-laws.

The extreme variability makes it hard to determine whether the size and duration exponents match other data, but the fitted scaling relation exponent was much less variable and more clearly separated from MEA or V_m_ results. The matched difference of median test (Wilcoxon signed-rank) between 49 recording groups found that the best fit *τ*(*τ* = 1.90 ± 0.63) was not significantly distinguishable from the V_m_ data (*r_SDF_* = 0.15,*p* = 0.33), but *β*(*β* = 1.41 ± 0.9) was dissimilar with a similar effect size (*r_SDF_* = 0.17,*p* = 0.028), and *γ_f_*(*γ_f_* = 1.11 ± 0.02) was also dissimilar (*r_SDF_* = 0.25,*p* = 7.1 × 10^−15^).

When comparing to the 13 samples of MEA data *γ_f_* was significantly different from the MEA data (*r_SDF_* = 0.88, and *p* = 9.21 × 10^−08^). This contrasts with our comparison between V_m_ and MEA data. In that case the scaling relation was not distinguishable even with 51 points of comparison and very low variability making a difference easier to detect. However, because of their extreme variability the size and duration exponents fail a 5% significance threshold for distinguishing from the MEA data by a Wilcoxon rank-sum result (*r_SDF_* = 0.06, *p* = 0.766 for *τ*and *r_SDF_* = 0.29,*p* = 0.123 for β). This failure of inverted LFP to show the same statistical properties as multi-unit activity may add a caveat to the assumptions behind the use of inverted LFP as a proxy for population activity (Beggs and Plenz, 2003; Kelly et al., 2010; Einevoll et al., 2013; Okun et al., 2015).

To summarize, the single-site LFP fluctuation results from the superposition of local spiking and extracellular synaptic current of juxtaposed network elements (Kajikawa and Schroeder, 2011; Einevoll et al., 2013; Pettersen et al., 2014; Ness et al., 2016). These fluctuations were found to be less informative about the network dynamics than single-neuron V_m_ fluctuations. V_m_ fluctuations result from the superposition of EPSPs and IPSPs indicating neuronal responses propagating in a manner consistent with the true neural network architecture. In other words, synaptic and spiking events driving fluctuations at single extracellular electrodes may be too badly out of sequence and distorted to faithfully represent neuronal avalanches, whereas the sequence of synaptic and spiking events driving somatic membrane potential fluctuations is functionally relevant by definition.

### Stochastic surrogates are distinguishable from V_m_ or MEA results, reveal importance of non-linear filtering

After eliminating inverted LFP as an alternative single-electrode signal, it was important to establish whether our results could have been created from a linear combination of independent random processes (Touboul and Destexhe, 2017; Priesemann and Shriki, 2018), similar to those used when contesting evidence for critical brain dynamics (Bédard et al., 2006; Touboul and Destexhe, 2010; Touboul and Destexhe, 2017). We also wanted to learn what effects non-linearity (non-Gaussianity) has in signals like the V_m_.

To address these questions, we used both the AAFT and UFT phase shuffling algorithms (see Methods: Experimental Design and Statistical Analysis). AAFT (**Figure 6**) preserves both the exact power-spectrum (autocorrelation) of the signal and non-linear skew of signal values but randomizes the phase (higher-order temporal correlations). UFT is the same but forces the distribution of signal values to be Gaussian. Using both allows us to attribute some characteristics to non-linear rescaling and others to precise temporal correlation structure.

Phase shuffling tends to preserve power-laws since it explicitly preserves the 1/f trend of the power-spectrum. However, the matched signed-rank test reveals that the values of the exponents change in both methods. Under UFT transformation the scaling relation and shape collapse became more trivial and like the LFP. This suggests that both the non-linear rescaling of input currents by membrane properties and the way that input populations interact throughout the intricate dendritic arborization are important.

For the 51 recording groups from the first 20-minutes the AAFT reshuffled data yield a median size exponent of *τ* = 1.74 ± 0.29 while the duration exponent was *β* = 2.0 ± 0.34 (**Figure 7D**). The fitted scaling relation exponent was *γ_f_* = 1.19 ± 0.06 and the predicted scaling relation exponent was *γ_p_* = 1.21 ± 0.49.

Pairing the surrogates to the original V_m_ data and performing the Wilcoxon signed-rank test for difference of medians gives (*r_SDF_ =* 0.053, *p* = 2× 10^−4^), (*r_SDF_* = 0.091, *p* = 0.08), and (*r_SDF_* = 0.207,*p* = 3 × 10^−5^) for *τ,β*, and *γ_f_* respectively. Thus *τ* and *γ_f_* are both significantly different, this is supported by the fact that only 55% of the groups meet all four standard criteria for criticality, while 76% of meet them for the original V_m_ time series. This difference between success rates is significant by Fisher’s exact test (*r_OR_* = 2.67,*p* = 0.0363).

The failure mode for AAFT shuffled data was almost entirely in reduced goodness of fit (*R*^2^) for a power-law fit to its scaling relation, 17% fewer recording groups met the criterion *R*^2^ > 0.95, than for V_m_ (*r_OR_* = 4.18,*p* = 0.0093). When the shape collapse is examined, we see another clear, if qualitative, difference in the symmetry of any presumed scaling function (**Figure 6C**). When taken together can we see that the AAFT shuffled dataset is not consistent with critical point behavior. Thus, we show that the exponent values and evidence for criticality, especially scaling and shape collapse which we inferred from V_m_ are not likely to come from random processes and are dependent on non-linear temporal correlation structure.

The key feature of the UFT result is that the fitted scaling relation exponent is much lower, *γ_f_* = 1.05 ± 0.049, which is significantly less than for AAFT (*r_SDF_* = 0.25, *p* = 1× 10^−13^) and less than the LFP (*r_SDF_* = 0.228,*p* = 3 × 10^−6^). It is very close to trivial scaling but is still distinguishable from *γ_f_* =1 at a population level via the sign test (*r_SDF_* = 0.843,*p* = 2× 10^−10^). Because the fitted scaling relation exponent and shape collapse were similar in both the UFT and LFP data, it suggests that lack of non-linear rescaling (non-linear filtering) may be a key feature of LFP that explains its failure to accurately reflect critical point behavior.

The UFT was universally poorer performing, 39% do pass the criticality test but given that the scaling relation exponent is so low this is simply random chance, and significantly worse than the V_m_ results (*r_OR_* = 5.04,*p* = 3 × 10^−4^). The UFT phase shuffling results obtain a median size exponent of *τ* = 1.69 ± 0.45 while the duration exponent was *β* = 1.81 ± 0.49. The predicted scaling relation exponent was = 1.01 ± 0.72. All are significantly different from the V_m_ results (*r_SDF_* = 0.183, *p* = 0.005), (*r_SDF_* = 0.199, *p*=2× 10^−4^), and (*r_SDF_* = 0.249,*p* = 2 × 10^−13^) for *τ,β*, and *γ_f_* respectively. These results are redundant with the AAFT confirming that our results do not have a trivial explanation.

When the scaling relation was examined, we saw another clear, if qualitative, difference in the symmetry of any presumed scaling function (**Figure 6C**). When taken together, our four standardized criteria followed by shape-collapse analysis let us distinguish phase-shuffled V_m_ fluctuations from the original V_m_ fluctuations, even limiting ourselves to data that meets the four criteria. Thus, the phase-shuffled data showed that the evidence for criticality in the original V_m_ fluctuations are dependent on non-linear temporal correlations.

### Excitatory and Inhibitory Synaptic Activity are Both Required for V_m_ Fluctuations to Match MEA Avalanches

Having learned that single-site LFP recordings cannot be used to accurately infer the statistics of population activity, and knowing that low-pass filtered and inverted LFP is believed to reflect *excitatory* synaptic activity (Kajikawa and Schroeder, 2011; Buzsaki et al., 2012; Einevoll et al., 2013; Ness et al., 2016) it begs the question: to what extent do excitatory synaptic events contain evidence for network criticality?

Somatic V_m_ fluctuations are the complex result of spatially and temporally distributed excitatory and inhibitory synaptic inputs further mangled by active and passive membrane properties in dendrites and soma. There is reason to believe that these features conspire to enforce the condition that V_m_ faithfully represents inputs to the presynaptic network (Boerlin et al., 2013; Barrett et al., 2014; Denève and Machens, 2016) similar to how input signals relate to presynaptic populations in our model. To address the stated question, we estimated the excitatory synaptic conductance changes 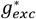 from the V_m_ recordings, using a previously developed inverse modeling algorithm (Yaşar et al., 2016), and applied the avalanche analysis on the inferred 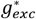 time series, (**Figure 6**).

The inferred excitatory conductance is plausibly related to the presynaptic population, however it failed to be a reliable measure of network dynamics (**Figure 7B**). We can’t know whether the failure is because excitatory current does not contain enough information or because the signal’s time constant is too short. Power laws in the avalanche size and duration distributions were observed in only 12% of the 51 groups from the first 20 minutes of recording. Comparing to V_m_ this was very different (*r_OR_* = 375,*p* = 6 × 10^−14^). Shape collapse was absent from the inferred excitatory conductance (**Figure 6C**) and none passed all four criteria for criticality. From this we conclude that inferred excitatory conductances are not a good network measure.

One of many potential reasons for this failure could be the much shorter time constant of the inferred 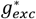 signal compared to the V_m_ signal. We saw exactly that situation when examining model results: *P_i_*(*t*) failed to reproduce network values as well as its smoothed version *ϕ_i_*(*t*). Therefore, we smoothed the 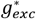 signal with an alpha-function, chosen because it should impose a similar non-Gaussian distribution as the V_m_ signal. The time constant of the alpha function was tuned to minimize the error between the autocorrelation of the smoothed 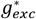 signal and the original V_m_ signal. By doing so we create a signal with a 1/*f* power-spectrum that should exhibit power-laws and reproduce many V_m_ statistical features, (**Figure 6**).

Reinstating the autocorrelation does not summon the return of scale-freeness. The smoothed signal did demonstrate power-laws (94%) and one serendipitously met the standardized criteria for consistency with critical point behavior (**Figure 6D**). However, this is chance. The average coefficient of determination for a fitted scaling relation on a log-log plot was *R*^2^ = 0.84 ± 0.14 so overall average avalanche sizes did not scale with duration as a power-law. Nonetheless this is a substantial improvement on the unsmoothed version *R*^2^ = 0.68 ± 0.17. This is a statistically significant difference (*r_SDF_* = 0.054, *p* = 3 × 10^−4^).

The smoothed inferred 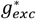 signal (**Figure 6A**) is visually more like the original V_m_ (**Figure 2B**) than the AAFT shuffled V_m_ surrogate (Figure 6A), however, it was a worse match. This shows that signals dependent only on excitation, even ones with the same non-Gaussian distribution and power-spectrum trend do not reflect the statistics of population activity. Interactions between EPSPs and IPSPs may be needed.

In conclusion, the single-site local field potential (LFP), the phase-shuffled recorded V_m_, and the inferred excitatory conductance 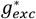, including its smoothed version, all failed to reveal the critical network dynamics. However, there are either similarities between the signals or some remaining scale-free signatures which reveal the importance of signal aspects. In order to faithfully represent population activity statistics a candidate signal must: have the right non-Gaussian distribution, the right 1/*f* power-spectrum characteristics and is sensitively dependent on higher-order temporal correlations such as may result from the complex interplay of excitation and inhibition within the dendritic arborization of a pyramidal neuron in the visual cortex.

## Discussion

Leveraging membrane potential (V_m_) and local field potential (LFP) recordings with modeling and microelectrode array (MEA) data yielded two principle findings: subthreshold V_m_ are a superior indicator of network activity and this correspondence is inherent to critical branching. Scrutinization revealed that avalanche size and duration distribution parameters covary to maintain stable geometrical scaling across different experiments, a noteworthy observation. The following discussion emphasizes possible significance and research intersections, such as explaining disagreement with theory via subsampling effects or quasicriticality or attempts to relate neural computation to the mathematical apparatus predicting behaviors of critical systems.

While “appropriating the brain’s own subsampling method” is a novel description of whole-cell recordings, it was inspired by examples. Whole-cell recordings contain information about the network (Gasparini and Magee, 2006; Mokeichev et al., 2007; Poulet and Petersen, 2008; El Boustani et al., 2009; Okun et al., 2015; Malina et al., 2016; Hulse et al., 2017; Lee and Brecht, 2018) and stimulus (Anderson et al., 2000; Sachidhanandam et al., 2013). Usually the focus is using neural inputs to predict outputs, not measuring population dynamics (Destexhe and Paré, 1999; Carandini and Ferster, 2000; Isaacson and Scanziani, 2011; Okun et al., 2015). Additionally, long-time or large-population statistics, like our adapted avalanche analysis, are useful for understanding neural code (Sachdev et al., 2004; Churchland et al., 2010; Crochet et al., 2011; Graupner and Reyes, 2013; McGinley et al., 2015; Gao et al., 2016) and are robust to noise. Our finding that single V_m_ recordings reflect scale-free network activity is significant as recording stability in behaving animals improves (Poulet and Petersen, 2008; Kodandaramaiah et al., 2012; Lee and Brecht, 2018). We open the door to using V_m_ fluctuations as windows into network dynamics.

Rigorous analysis supports our main experimental finding: subthreshold V_m_ fluctuations mimic neuronal avalanches and evince critical branching but negative LFP deflections don’t, despite being purported network indicators (Bédard et al., 2006; Liu and Newsome, 2006; Kelly et al., 2010; Einevoll et al., 2013; Okun et al., 2015). Our results and interpretations originate from spontaneous activity of ex-vivo turtle visual cortex. However, it is reasonable to hypothesize that some results generalize, because the turtle visual cortex shares much with other species, including recurrent cortical circuits bestowing pyramidal neurons with receptive fields and response tuning, although retinotopy is absent (Ulinski, 1990; Larkum et al., 2008). Lastly, the results are not coincidental measurement of noise because we could distinguish the V_m_ dataset from a surrogate dataset with identical power-spectrum and signal envelope but randomized phase (Theiler, 1992).

Readers keen on critical branching may notice our exponents differ from the exact theoretical predictions (*τ* = 1.5, *β* = 2 (Haldeman and Beggs, 2005)). Others observing this mismatch have suggested the brain operates slightly off-critical (Hahn et al., 2010; Priesemann et al., 2014; Tomen et al., 2014).

An extension of this suggestion, quasicriticality (Williams-Garcia et al., 2014), also explains the highly stable scaling relation: biological systems blocked from precise critically may optimize properties which are maximized only for critical systems, becoming “quasicritical”. Correlation time and length are maximized only at criticality and closely related to avalanche geometrical scaling (Tang and Bak, 1988; Sethna et al., 2001). If brains optimize correlation length, a highly stable scaling relation may result. Furthermore, including inhibition (Larremore et al., 2014) makes our otherwise critical model less consistent with criticality except that population statistics can still be inferred from input fluctuations. The stable-scaling was not in the model, which lacks any plasticity mechanisms. Stable-scaling may be a rare observation of self-organization principles such as quasicriticality. A contributing explanation is subsampling effects (Priesemann et al., 2009; Levina and Priesemann, 2017) but it doesn’t explain the stable scaling relation unless quasicriticality is also invoked.

### Neuronal Avalanches and Neural Input Fluctuations Similarity is Captured by a Critical Recurrent Coarse-Graining Network

Our main modeling finding, inputs to a neuron reflect network activity best for critical branching networks, is supported by a parameter sweep and detailed analysis. The implications are transferrable to networks where neural inputs fluctuate about proportionality to some subsample’s activity. We tune proportionality to be one, but that can also emerge from plasticity (Shew et al., 2015; Del Papa et al., 2017). Tight-balance suggests a biological mechanism causing subthreshold V_m_ to track excitation into a presynaptic population because IPSPs can have their timing and strength “balanced” to truncate EPSPs which would otherwise last longer than spurts of presynaptic excitation (Boerlin et al., 2013; Barrett et al., 2014; Gatys et al., 2015; Denève and Machens, 2016). We use V_m_ proxy, *ϕ_i_*(*t*), an alpha function convolved with a point process, *P_i_*(*t*). This *ϕ_i_*(*t*), is more like V_m_ than *P_i_*(*t*) and reproduces our experimental findings. Lastly, we investigate quasicriticality by including inhibition but tuning the maximum eigenvalue to what would be the critical point without inhibition.

Our model provides insights on network subsampling and renormalization group. Usually subsampling means selecting neurons at random or modeling an MEA with an arbitrary grid (Priesemann et al., 2009). Our “subsample” is the presynaptic population represented by summing weighted inputs from active neurons. This is the first analysis intersecting actual convergence of network flow (i.e. postsynaptic soma).

Subsampling effects distort avalanche size and duration distributions, likely creating differences between experimental results and theoretical predictions (Priesemann et al., 2009; Ribeiro et al., 2014; Levina and Priesemann, 2017). Subsampling may explain the small disagreement between avalanche analysis performed on simulated network activity, *F*(*t*), V_m_ proxy *ϕ_i_*(*t*), and single-neuron firing rate *P_i_*(*t*). However, V_m_ and MEA results are off theory but match each other. Either their subsampling errors are alike enough to produce similar distortions, or subsampling co-occurs with some form of quasicriticality (Priesemann et al., 2014; Williams-Garcia et al., 2014).

Intriguingly, the Restricted Boltzmann Machine (RBM) (Aggarwal, 2018), (a related model) was exactly mapped to a “renormalization group” (RG) operator (Mehta and Schwab, 2014; Koch-Janusz and Ringel, 2018). RG is a mathematical apparatus relating bulk properties to minute interactions (Maris and Kadanoff, 1978; Nishimori and Ortiz, 2011; Sfondrini, 2012). It characterizes critical points of phase-transitions (Stanley, 1999; Sethna et al., 2001) and helps derive neuronal avalanche analysis predictions (Sethna et al., 2001; Le Doussal and Wiese, 2009; Papanikolaou et al., 2011; Cowan et al., 2013). RG operators have a coarse graining step and a rescaling step, reminiscent of resizing a digital image. Crucially, iterating the appropriate operator on a critical system produces statistically identical “copies”, but on non-critical systems the iterations diverge. Our experimental findings are reproduced by a model which averages (coarse grains) presynaptic pools to get an instantaneous firing probability for each neuron. Then a logical operation (rescaling) sets the spiking states for the next iteration, demonstrating an RG-like operation. Denève and Machens (2016) proposed a similar relationship between real V_m_ and presynaptic pools. The finding that RBMs converge to a similar neural operation underscores the relevance of RG. The importance is that a recurrent network of RG operators may be like a criticality ouroboros, displaying widespread scale-free signatures if any component is critical or briefly driven by critical or scale-free inputs (Mehta and Schwab, 2014; Schwab et al., 2014; Aoki and Kobayashi, 2016; Koch-Janusz and Ringel, 2018).

Significantly, associating neuronal processing with critical branching may induce an organizing principle, the “Information Bottleneck Principle”. This foundational concept for deep learning balances dimensionality reduction (compression) against information loss (Tishby and Zaslavsky, 2015) and is reminiscent of both efficient coding (Friston, 2010; Denève and Machens, 2016), and origins of tuning curves (Dayan and Abbott, 2001; Wilson et al., 2016; Heeger, 2017). Koch-Janusz and Ringel (2018) trained their network by maximizing mutual information between many inputs and fewer outputs. This produced nodes with receptive fields matching the coarse graining step of popular RG operators. They derived correct power-laws by iterating the network. Applications of RG to neural computation are widespread: image processing (Gidas, 1989; Mehta and Schwab, 2014; Saremi and Sejnowski, 2016), brain and behavior (Freeman and Cao, 2008), emergent consciousness (Werner, 2012; Fingelkurts et al., 2013; Laughlin, 2014), and hierarchical modular networks (Lee et al., 1986; Willcox, 1991) important for criticality (Moretti and Munoz, 2013). Furthermore, our model’s RG-like features are crucial to reproducing experimental results. It follows that elegant RG operators like in the RBM might also capture biological neuronal processing, fulfilling the demand for beautiful neuroscience models (Roberts, 2018) while offering insights into organizing principles and observations of scale-freeness.

## Conclusion

We established that subthreshold fluctuations of V_m_ in single neurons agree with neuronal avalanche statistics and with critical branching but fluctuations in other single-electrode signals do not. Computational modeling showed that accurate inference requires critical branching like connectivity. Fluctuation size scales with duration more self-consistently in experimental than model results, hinting at self-organization. These findings are consistent with a nascent reduction of neural computation to coarse-graining operations which may explain the prevalence of critical-like behavior during spontaneous neural activity. Fully articulating the implications requires more investigation, but we have substantially extended the evidence for critical branching in neural systems while rigorously demonstrating that subthreshold V_m_ fluctuations of single neurons contain useful information about dynamical network properties.

## Data and Software Accessibility

All raw data and the code developed for this analysis is available upon request to the corresponding author: James Kenneth Johnson.

## Acknowledgements

Woodrow Shew at the University of Arkansas for helping with the comparison to MEA data. Nathaniel (Caleb) Wright is now at Georgia Tech.

This research was supported by a Whitehall Foundation grant (no. 20121221; R. Wessel) and a National Science Foundation Collaborative Research in Computational Neuroscience grant (no. 1308159; R. Wessel).

